# MAMBA: a model-driven, constraint-based multiomic integration method

**DOI:** 10.1101/2022.10.09.511458

**Authors:** Manuel Ugidos, Carme Nuño-Cabanes, Sonia Tarazona, Alberto Ferrer, Lars Keld Nielsen, Susana Rodríguez-Navarro, Igor Marín de Mas, Ana Conesa

**Affiliations:** Gene Expression and RNA Metabolism Laboratory, Instituto de Biomedicina de Valencia, Consejo Superior de Investigaciones Científicas, Valencia, Spain; Department of Applied Statistics, Operations Research and Quality, Universitat Politècnica de València (UPV), Valencia, Spain; Novo Nordisk Foundation Center for Biosustainability, Technical University of Denmark, Copenhagen, Denmark; Genomics of Gene Expression Laboratory, Institute for Integrative Systems Biology, Spanish National Research Council, Valencia, Spain

**Author notes:** These authors jointly supervised this work (Igor Marín de Mas, Ana Conesa).

## Abstract

The inclusion of omic data into constraint-based modeling (CBM) has improved metabolic network characterization of biological models. However, the integration of semi-quantitative metabolomic data into CBM remains challenging. Here, we present MAMBA (Metabolic Adjustment via Multiomic Blocks Aggregation), a CBM approach that enables the use of semi-quantitative metabolomic data together with a gene-centric omic data type (e.g. transcriptomics, ChIP-seq or chromatin accessibility data, among others), and the combination of different time points and conditions. MAMBA outperformed other methods in terms of metabolic network characterization and metabolite prediction accuracy. As a case study, we applied MAMBA to a yeast multiomic dataset with time series design where two different yeast strains where exposed to heat stress. MAMBA captured known biology of heat stress in yeast and identified novel affected metabolic pathways. MAMBA was implemented as an integer linear programming (ILP) problem to guarantee efficient computation, and coded for MATLAB (freely available at github.com/ConesaLab/MAMBA).

## 1 Introduction

The availability of sequenced genomes, together with the annotation of genes and their functions has facilitated the reconstruction of high-quality genome-wide metabolic networks, the so-called, genome-scale metabolic models or GEMs [1]. These metabolic networks gather all the known metabolic reactions identified in an organism’s genome and incorporate information about stoichiometry, thermodynamics and optionally the associations between the reactions and proteins/genes involved. Constraints-based modeling (CBM) is a family of mathematical methods suitable for the analysis of these large metabolic networks. When CBM is applied to GEMs, the fluxes through metabolic reactions represent the model variables to be estimated. The computation of each variable in CBMs is constrained by a minimum and maximum range of values. CBMs calculate flux distribution through the metabolic network that satisfies two fundamental types of constraints [2] i) steady-state mass-balance, which sets the total production and consumption rates for each metabolite to be equal; ii) capacity, i.e., upper and lower bounds for fluxes can be imposed. One of the most widely used CBMs is Flux Balance Analysis (FBA). This method describes the phenotype of an organism making use of an objective function (OF) that needs to be optimized (i.e. biomass maximization) [3]. The OF, together with the stoichiometric and thermodynamic parameters -embedded in the metabolic model- and the imposed constraints are formalized as numerical matrices that can be solved by a number of mathematical optimization algorithms and software developed for this purpose to define tissue/organism-specific metabolic network flux profile [4].

In order to improve the metabolic network characterization by CBM methods, additional biological data can be included into GEMs. One of the first extended CBM method was the GIMME algorithm [5], that proposed the incorporation of gene expression data into the FBA model. GIMME connects a given metabolic reaction to the genes encoding the enzymes that carry out the reaction. To determine the output of reaction fluxes, GIMME minimizes the usage of reactions where lowly-expressed genes participate, while keeping the OF above a certain value. Reactions and genes are connected by a set of Boolean rules that associate reactions to the expression state of the involved genes using a binary representation. These rules are known as gene-protein-reaction associations or GPRs and they indicate the collection of proteins (isozymes, enzymatic subunits, etc.) required for the reaction to carry flux. To determine the expression state of genes, GIMME and other CBMs require a set of user-supplied expression thresholds for classifying genes, and in turn reactions, generally as on or off [6], although more states can also be used. These arbitrary thresholds have a strong impact in the output of the method and hence must be carefully selected, which is an important caveat of such CBMs. Notably, other approaches that do not require user-supplied expression threshold values have also been developed, e.g. MADE [7]. MADE relies on expression data from two or more experimental conditions and uses the results of a differential expression analysis to determine gene states across conditions. The inclusion of transcriptomics data into CBMs has been successfully used for predicting associations between silencing or expression of genes and the metabolic capabilities of an organism [1, 8]. These approaches rely on the proven existence of a correlation between transcript levels and reaction activity [9] and it has been widely demonstrated that GEMs become more powerful by integrating additional molecular information [10].

Metabolomic data have also been included into CBMs to improve metabolic network characterization [11–14]. Metabolomics are typically integrated into GEM reconstruction analyses as a set of capacity constraints that limit the flux through a given reaction/s. To this aim, metabolomic data from experiments using isotopically labeled subtrates or absolute quantification from label-free experiments can be used [15]. However, the inclusion of semi-quantitative (relative quantification) metabolomics into CBM still remains challenging. Furthermore, there is growing evidence of a significant contribution of epigenomics features such as chromatin modifications to the regulation of the metabolism and vice versa [16–18]. However, the incorporation of chromatin modification data into the CBM framework has not been yet reported. Finally, to the best of our knowledge, methods enabling the integration of non-quantitative metabolomics and transcriptomics data combining different time points and conditions are still not available. All together, these considerations imply that despite the power of CBM for modelling metabolic fluxes, there is a significant number of applications for which these methods have not been adapted yet. This represents a gap in our understanding of metabolic regulation at large, and a missed opportunity for the analysis of a wealth of multi-omics data that include metabolomics measurements together with other molecular layers and/or a variety of experimental designs.

In this work we present MAMBA (Metabolic Adjustment via Multi-omic Block Aggregation), a constraint-based genome-scale metabolic reconstruction model that integrates gene-centric omics measurements such as gene expression or histone modifications without requiring arbitrary expression thresholds, and semi-quantitative metabolomic data. Any experimental design can be modeled with MAMBA. When analyzing time-series data, MAMBA considers the dynamics of the system by including all time-points in the same model. MAMBA has been tested with a multi-omic time-series yeast dataset consisting of two strains, *mip*6Δ mutant and Wild Type (WT) subject to heat-shock treatment. Samples were obtained at three different time points: baseline, 20 min. and 120 min., after heat-shock and profiled for metabolomics, RNA-seq and ChIP-seq. MAMBA showed a better performance than other CBMs to predict metabolite changes, found key differences between strains regarding dynamic adaptation to heat stress and revealed differences between the transcriptional and chromatin control of metabolic fluxes. MAMBA was implemented in MATLAB and it is freely available at https://github.com/ConesaLab/MAMBA.git.

## 2 Data and computational details

In this work we have used a multiomic dataset from yeast previously generated in our lab and detailed at Nuno-Cabanes et al., 2020 [19]. Briefly, this dataset consists of gene expression RNA-seq, NMR targeted metabolomic data and H4K12ac ChIP-seq data generated from the same samples. Yeast cells were subjected to heat stress and measurements were taken at three different conditions: 30 ° C 20 min., 39 ° C 20 min. and 39 ° C 120 min. Two different strains were analyzed: Wild Type (WT) yeast cells and a strain lacking the mip6 gene (*mip*6Δ), which is a factor involved in mRNA metabolism [20]. The improved iMM904 *Saccharomyces cerevisiae* GEM model [21] was used as the yeast metabolic network. This metabolic model contains 1226 metabolites, 1577 reactions and 905 genes. RNA-seq data covered the totality of metabolic genes in the GEM, however only 30 metabolites of the metabolic model were present in the NMR data, that included 4 sugars, 12 amino-acids, 4 alcohols, 5 carboxylic acids, and other compounds (CMP, NAD, Glutathione, ATP and GMP).

RNA-seq, ChIP-seq and metabolomics data were pre-processed as described in Nuno-Cabanes et al., 2020 [19]. Differential gene expression and H4K12ac levels were calculated using the limma package [22] and FDR corrected p-values were used. The MAMBA method was implemented in Matlab, and Gurobi library [23] was used as the default solver for linear programming problems.

## 3 Description of the approach

### 3.1 Flux Balance Analysis

Given a metabolic network model containing *M* metabolites and *L* reactions, FBA formalizes the mass balance of internal metabolites as a set of linear equations that satisfy the condition:

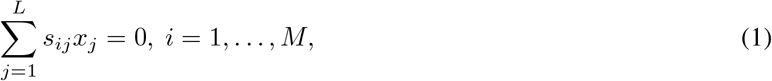

where *x*_*j*_ corresponds to the flux of reaction *j*, and *s*_*ij*_ stands for stoichiometric coefficient of metabolite *i* in reaction *j*. This condition represents the steady state or null neat mass balance, i.e. that for all cellular metabolites in the network the sum of all productions and consumptions equals zero. FBA calculates x (vector of fluxes for all reactions) given **S** (an *M* x *L* stoichiometric matrix), which is known from the metabolic network model. Capacity constraints can be added to the equation system as inequalities to delimit the solution space:

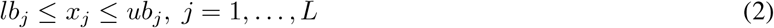

where *lb*_*j*_ contains the lower bounds of the reaction *j*, i.e., minimum flux value that reaction *j* is allowed to carry; and *ub*_*j*_ indicates upper bound of reaction *j*.

For most metabolic networks, this results in an undetermined system of equations. Thus, the FBA method makes use of linear programming (e.g.: mixed-integer linear programming or milp) to determine the optimal flux distributions given the objective function, OF (*f* (x)):

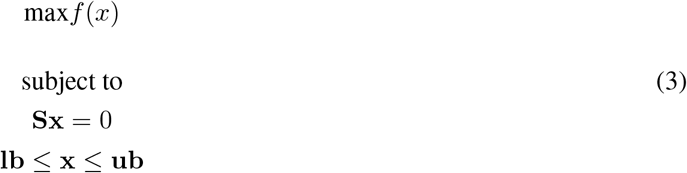

The OF, *f* (x), contains coefficients associated to the values of x to satisfy the restrictions imposed to the model. In regular FBA, the OF is usually the maximization of the biomass production reaction and therefore, only the coefficient of the reaction of biomass production biomass is set to be distinct from zero. However, the coefficients in *f* (x) can be customized to maximize any reaction in the metabolic network or combination of reactions.

Figure 1 describes FBA model construction for a small toy metabolic network, where biomass production is obtained through reaction *R*_5_ (*x*_5_). The metabolic network model provides all required objects except x, which is the model solution. FBA can only be applied to a single condition or biological status.

**Figure 1:**
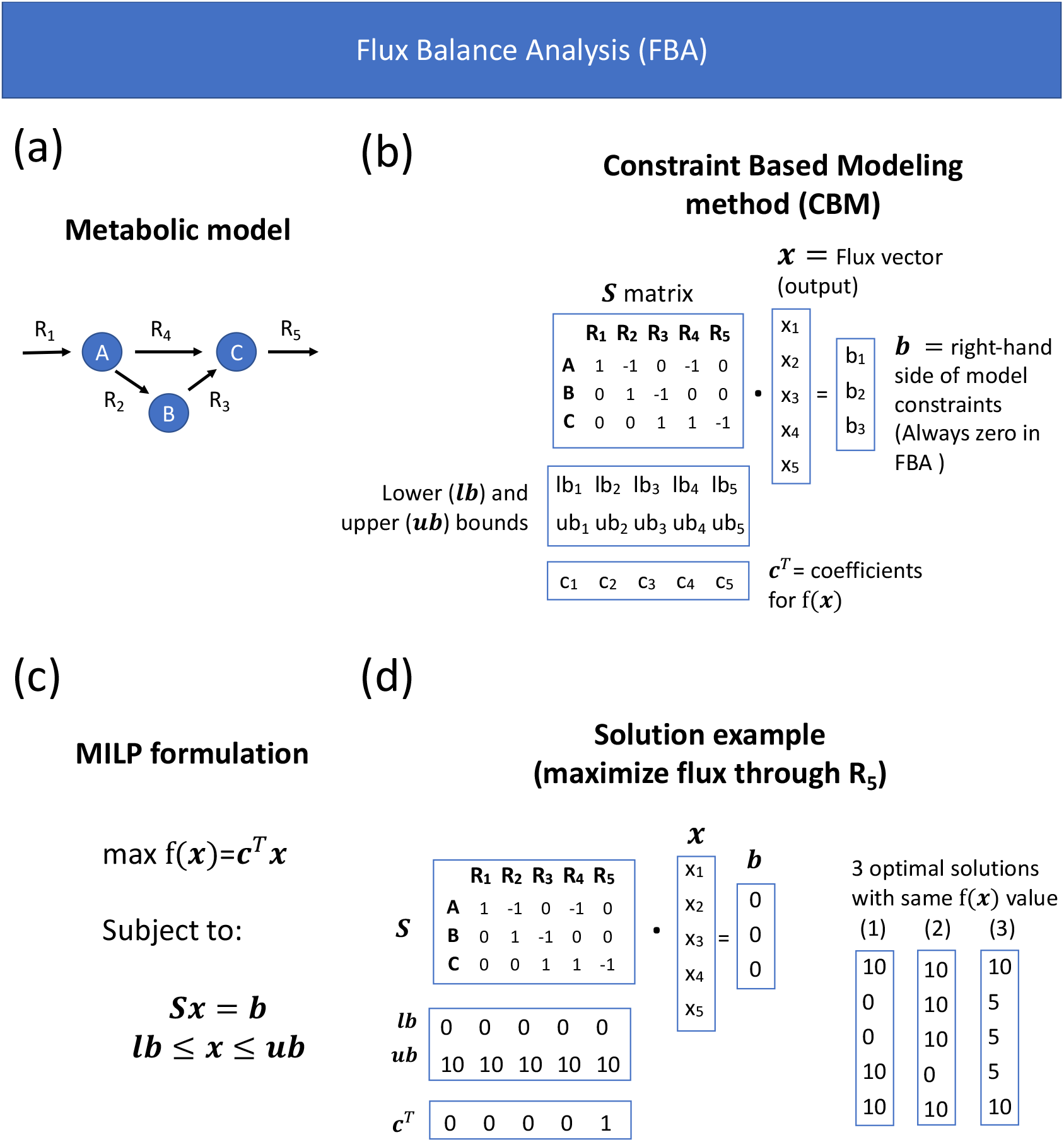
Illustration of FBA. a) Representation of the metabolic network. b) Mathematical representation of FBA elements. c) FBA formulated as a linear programming problem. d) Example of a FBA solution for the toy network.

### 3.2 Integration of gene/protein associated information into GEMs

Gene-protein-reaction rules (GPRs) describe the associations between genes/proteins and reactions by using Boolean definitions that represent the gene(s) encoding the protein(s) required to catalyze a given reaction. Therefore, GPRs unlock the integration of gene/protein associated omic data into metabolic models resulting into the so-called Genome-scale metabolic models or GEM, that contains both the metabolic network and the gene-reaction associations. GPRs are used to constraint the fluxes of metabolic reactions based on gene/protein-associated omic data. As shown in Figure 2, a metabolic reaction is allowed to be active if and only if the associated GPR is TRUE, which is determined by the expression state of the gene(s) involved. These GPR-dependent constraints are included within the capacity constraints into CBM. Additionally, MAMBA introduces stoichiometric GPRs (S-GPRs) into the model which provide the information about the sub-units resulting from the transcription of each gene required to have a catalytically active unit [24].

**Figure 2:**
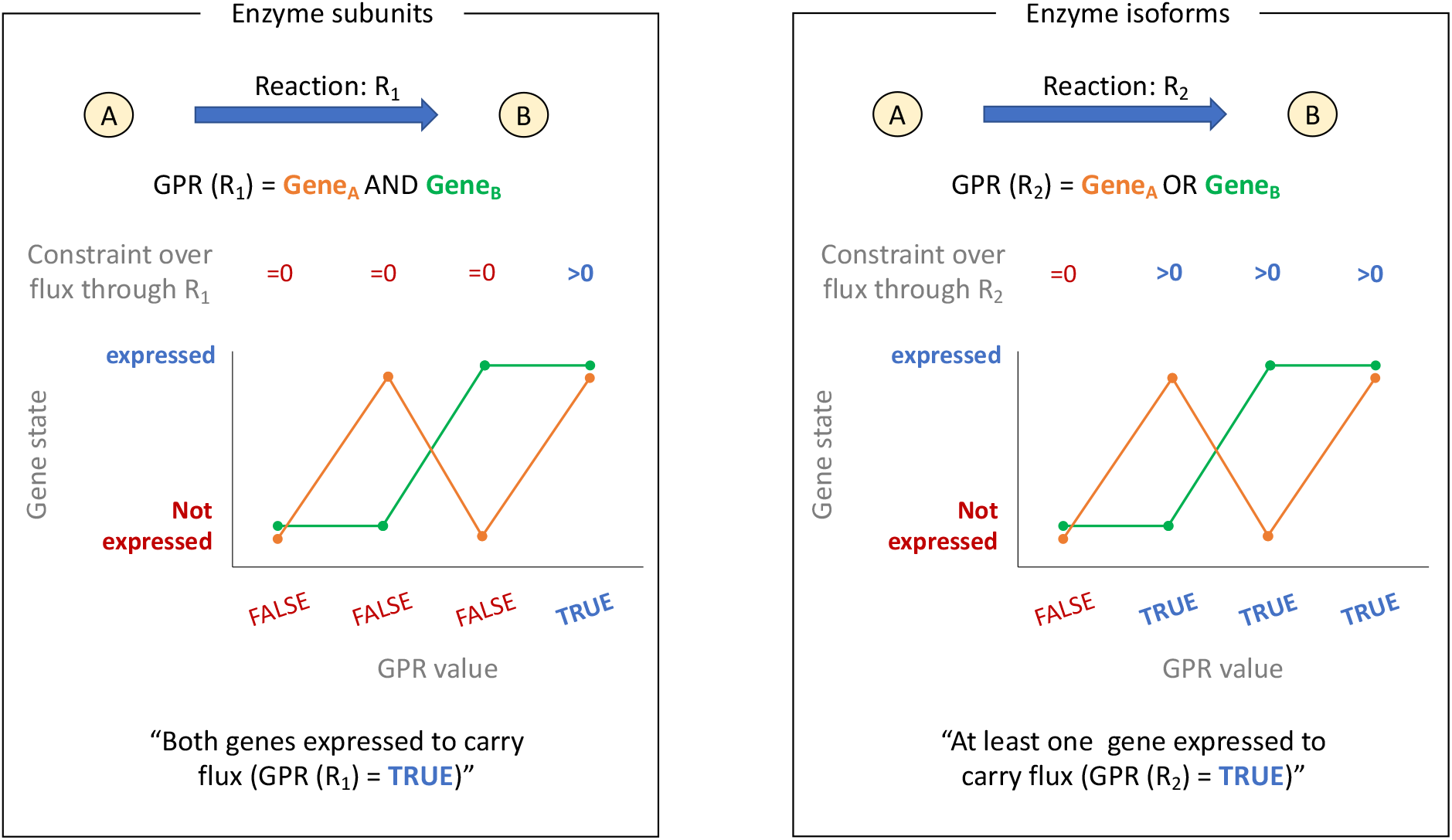
Graphical representation of GPRs. Left panel represents a GPR for enzyme subunits that have an “AND” relationship where both genes need to be expressed for the reaction to be active. Right panel represents a GPR for enzyme isoforms, which are modelled by “OR” relationships.

### 3.3 Formalizing multiomic-based constraints in MAMBA

MAMBA uses omic data on top of the metabolic model to constraint the FBA solution. In particular, a gene-centric omic data type (e.g. gene expression) and metabolomic data are used to formulate new model constraints. Therefore, MAMBA requires several input data in addition to the metabolic model. MAMBA works by simultaneously modeling all experimental conditions under study (two at least) as it uses relative feature quantification. Regarding gene-centric omic data, the algorithm needs the output of a differential feature analysis, i.e., an effect-size measure and the associated p-values. Similarly, metabolite ratios (or an effect-size measure as well) between conditions must be also given to the model as input data. MAMBA modifies the elements of FBA model (Equation 3) according to the input data as described below (Figure 3).

**Figure 3:**
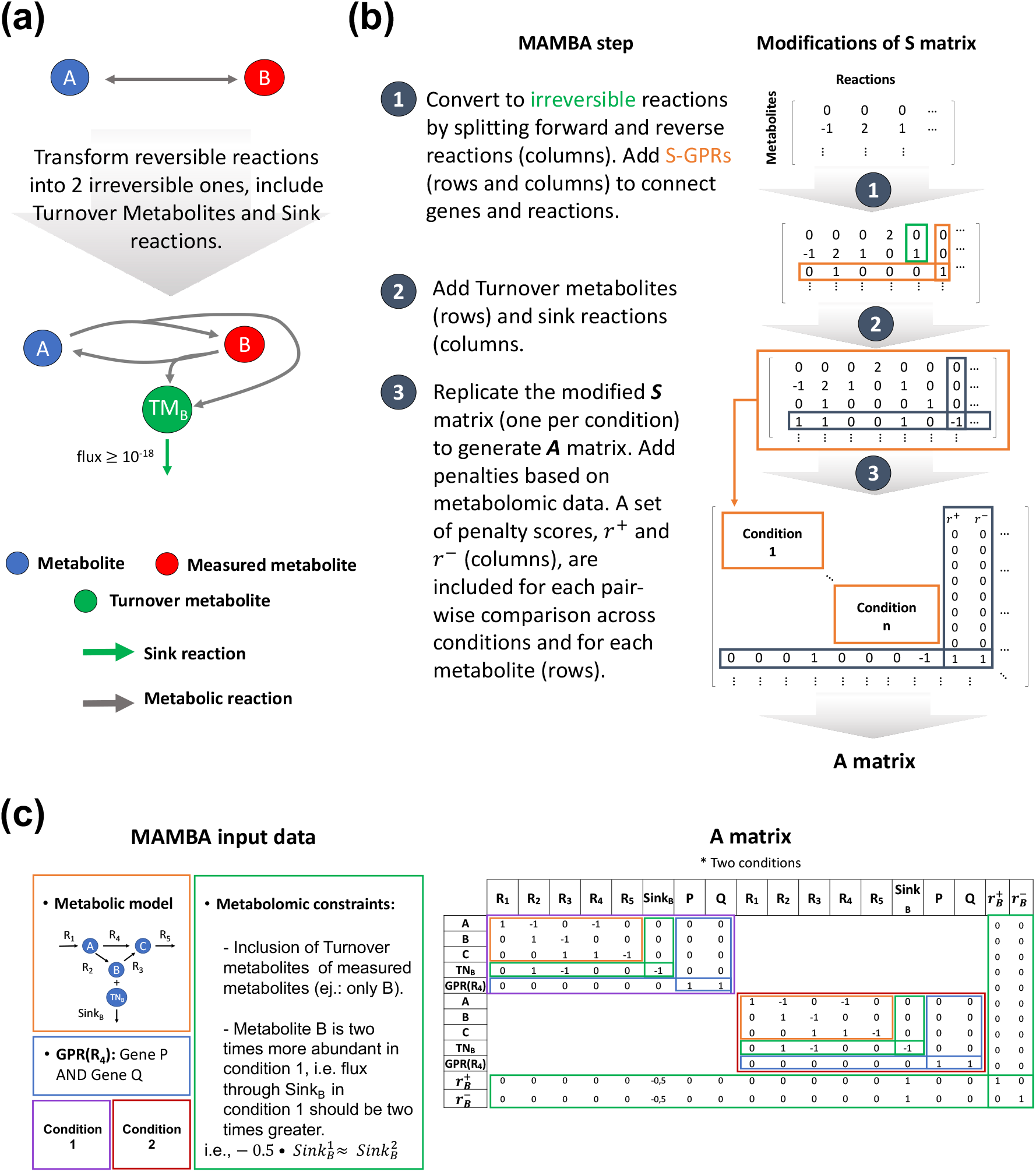
Graphical representation of model modifications performed by MAMBA. a) Modifications of the genome-scale metabolic model. b) MAMBA model is contained in **A** matrix resulting from the modification of the standard stoichiometric matrix (**S**). c) Structure of MAMBA model and input data require to construct **A** matrix.

#### 3.3.1 Gene/protein associated data

MAMBA works with differential values between two comparing conditions -the usual design of omics experiments-rather than with absolute values. By using differential data, MAMBA bypasses the difficult task of defining absolute threshold values to determine different levels of gene expression. More importantly, MAMBA allows the incorporation in the model of any type of omics data, as long as a gene/protein associated value can be computed, thereby unlocking the utilization of multi-omics data in FBA. In the reminder of the model formulation, we will use the notation gene to generally represent any omics measurement that can be associated to a gene ID such as expression, protein or chromatin data. Consequently, constraints over reactions can be generated based on different omic data types and, by comparing the resulting models, inferences can be done about the control of the metabolic network by different molecular regulatory layers. In this work, we demonstrate MAMBA using gene expression (RNA-seq) and histone modification (ChIP-seq) data, but other omics modalities, such as DNA methylation or chromatin accessibility could also be used. To incorporate gene-associated omics measurements, MAMBA adapts the MADE model [7] developed for gene expression data. The algorithm requires an effect size measure (typically, log2 Fold-change) or a statistic that compares gene values across conditions, together with the associated p-values. Based on the omics data, genes are classified into three categories: UP (gene activity increases significantly), DOWN (gene activity decreases) and CONSTANT (non-significant change). The algorithm finds a sequence of binary expression states (one gene state for each comparing condition) that best fits the differential expression data. Formally, the sequence of binary gene states returned by the algorithm is expressed as:

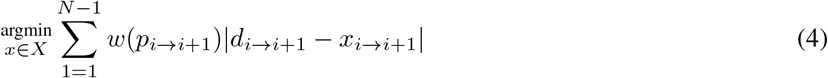

where *N* is the number of conditions, *I* → *i*+1 represents a transition from condition *n*_*i*_ to *n*_*i*+1_, x is the solution vector containing the predicted binary gene states, **p** contains the associated p-values for each transition, *w*() is a weighted function used to prioritize gene state calculations and **d** is the vector of observed differences being: 1 for UP genes, -1 for DOWN genes and 0 for CONSTANT genes. Typically the weighted function *w*() is the -log10(p-value) of the differential expression, implying that when two gene transitions (gene states) are not simultaneously feasible, p-values are used to prioritize and the transition associated to a lower p-value is reflected in the output of the model. MAMBA uses Equation 4 to find a solution of gene states (i.e., 0 or 1) that minimizes the differences between the observed and predicted gene state changes. However, in the formulation of the stoichiometric matrix, vector of differences (**d**) cannot be included and hence an equivalent expression of Equation 4 is used. Therefore, for a given transition *i* → *i* + 1, the OF for MAMBA can be defined as the following weighted sum:

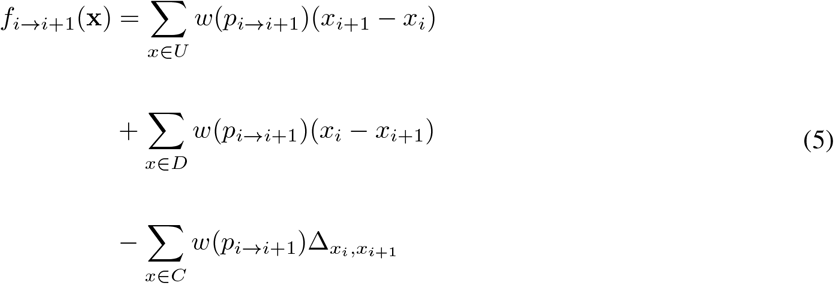

where U is the list of UP genes, D is the list of DOWN genes and C is the list of CONSTANT genes. Regarding UP genes, if the solution matches the expected output (*x*_*i*_ =0 and *x*_*i*+1_ = 1), the first element of this equation will be positive and therefore contributes to maximize *f* (x). The same occurs for DOWN genes as the position of *x*_*i*_ and *x*_*i*+1_ is swapped in the formula. Finally, the third element controls the contribution of constant genes to the OF by defining Δ _*xi,xi*+1_ as a binary variable that takes the value 0 when *x*_*i*_ = *x*_*i*+1_ and 1 otherwise. Thus, the third element is negative when the solution does not match the expected result, thereby penalizing *f* (x).

#### 3.3.2 Metabolomics data

Next to gene-associated omics data, MAMBA also incorporates into the model semi-quantitative metabolomics data obtained from comparing experimental conditions. To add metabolomics data to the MAMBA model, we incorporated the concept of sink reaction presented in Schmidt et al., 2013 [11]. This implementation consists of creating a new set of artificial metabolic reactions, the sink reactions, that connect measured metabolites to the GEM through a two-step process. First, reversible reactions are transformed into two irreversible reactions. Next, for each measured metabolite, a turnover metabolite is added to the model and connected to every reaction that produce or consume the actual measured metabolite. Finally, a sink reaction is included having the turnover metabolite as unique reactant and no products (Figure 3.a). To ensure that all detected metabolites have associated sink reactions with non-zero flux in the solution, the lower bound of sink reactions is set to a small positive value limited by the linear programming solvers numerical tolerance [10^−’8^ [23]].

Once turnover metabolites and their corresponding sink reactions are included in the model, constraints can be formulated. Basically, the observed metabolite ratios between conditions are modeled. For instance, let us consider a metabolite *m* measured in two conditions, *A* and *B* with values *m*_*A*_ =6 and *m*_*B*_ = 12. Since the quantification of *m* is two times greater in condition *B* than in *A*, the flux of *m* through the sink reaction in condition *B* should be twice the flux through the sink reaction in condition *A*. This ratio requirement can be added to the model by imposing both lower and upper bounds of sink reactions to have the same ratio relationship. However, such modification of the capacity (hard) constraints dramatically decreases the number of feasible solutions. Instead, a penalty on the OF, which is a soft model constraint, was implemented as the parameter *r*, which reflects the difference between the observed and the predicted ratio. In order to deal with positive and negative deviations of the solution with respect to the expected ratio, two different penalty scores are created, *r*^+^ and *r*^−^ (Figure 3.b and c). The difference between both is that *r*^+^ is constrained to be greater than or equal to zero and *r*^−^ is constrained to be lower than or equal to zero. To illustrate the meaning of these two penalty scores, let us consider *v* as the flux through the sink reaction associated to metabolite *m* and only two conditions (*A* and *B*), the penalty imposed is derived from:

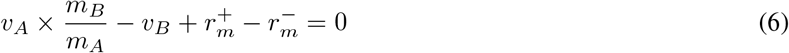

From the expression above, one can derive that the lower the values of 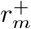 and 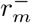, the higher the concordance between observed and modeled metabolite ratios. Therefore, the model is forced to minimize both penalty values:

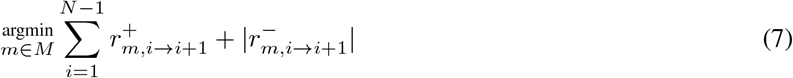

where *N* is the number of conditions and *M* is the number of metabolites (or number of sink reactions). Finally, the new component is included in the OF (Equation 5) and its final form is:

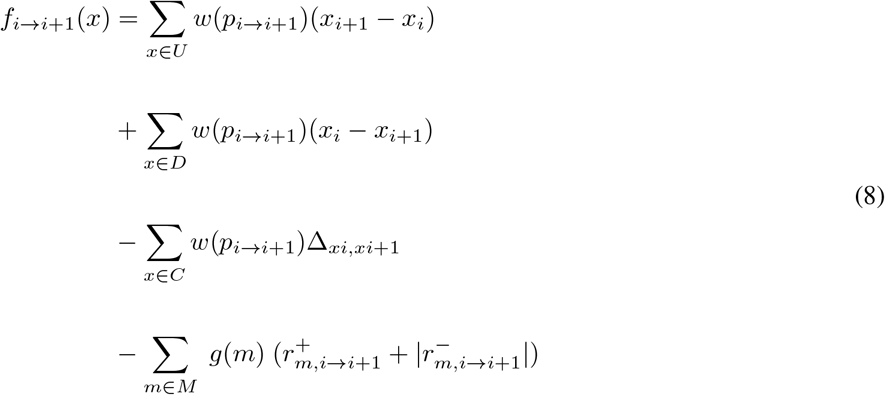

The expression above is the final OF that MAMBA maximizes to find the optimal solution for *i* = 1, …, *N*, being *N* the number of conditions. Note that an additional weight function for metabolomic constraints, *g*(*m*), has been included. This function has the purpose of balancing gene expression and metabolomics constraints to give them similar weights in the OF. Since constraints from gene expression include a weight function, *w*() based on p-values, these constraints may be very strong for very low p-values, e.g. *w*(1 × 10^− 100^) = 100 (considering a p-value = 1 × 10^− 100^). Additionally, without considering any weight function for metabolomic constraints, every deviation from the actual ratio has the same penalty. For instance, let us consider two metabolites *m*_1_ and *m*_2_ having observed ratios between 2 conditions *A* and *B* defined as: *m*_1,*A* → *B*_ = 3 and *m*_2,*A* → *B*_ = 6, while their predicted ratios are *m*_1, *Â* → *B*_ = 2 and *m*_2, *Â* → *B*_ = 5. Both metabolites would contribute to the OF with the same magnitude, 1, which is the penalty associated to the difference between observed and predicted ratios according to Equation 6. However, the prediction for metabolite *m*_2_ is more accurate than for metabolite *m*_1_ (5*/*6 *>* 2*/*3). To account for this issue, ratios of metabolites are standardized between 0 and 1, being 1 the highest observed ratio. The metabolite weight function, *g*(*m*) captures these two considerations by defining *g*(*m*) as:

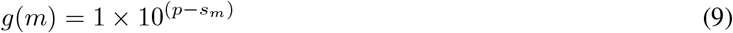

where *p* is the value of the highest p-value of the gene expression data after *w*() (− log10) transformation (considering only genes included into the GEM) and *s*_*m*_ is the standardized ratio of metabolite *m*. Considering the previous example with two metabolites, the standardized ratio for metabolite *m*_1_ is 0 and for *m*_2_ is 1. Hence, the penalty for *m*_1_ is *g*(*m*_1_)=1 × 10^(*p*−0)^, while the penalty for *m*_2_ is *g*(*m*_2_)=1 10^(*p* − 1)^, and therefore, penalty of *m*_2_ is lower than penalty of *m*_1_ (as *p* is a common value).

Finally, MAMBA representation as a linear programming optimization problem is defined as follows:

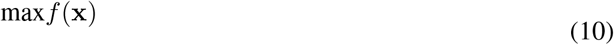

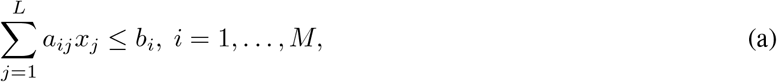

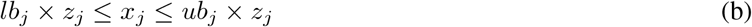

where **A** (with general element *a*_*ij*_) contains original stoichiometric matrix and transcriptomics and metabolomics constraints, vector **b** sets the boundaries of the constraints that can be different from zero, vector x is the optimal solution to be found and vector **z** contains the S-GPR constraints, i.e, *z*_*j*_ is 1 if the S-GPR is TRUE and 0 otherwise. Finally, the final OF including all transitions is represented as 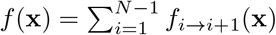

### 3.4 Evaluation of MAMBA method

#### 3.4.1 Sensitivity and Robustness analysis

The sensitivity analysis consists in evaluating the impact of each reaction state on the optimization function. Each reaction, one by one, was forced to be active (A) or inactive (I) at every condition by modifying capacity constraints, i.e. a given reaction is forced to be active by setting its lower bound *>* 0 and it is forced to be inactive by setting both, lower and upper bounds, equal to zero. The optimization function was evaluated in each case and compared to the MAMBA model without forced reactions. In case the result equals the unforced model, the reaction is deemed not to be part of the solution, while if the result is substantially different, the reaction is considered critical in the model. Therefore, sensitivity analysis identifies both the reactions that most affect the optimization function and the reactions with an undetermined state, i.e., the optimization result is the same whether they are active or inactive. Robustness analysis identifies the consistency of predictions as a function of changes in method parameters and identifies reaction states (Active or Inactive) that are unambiguously predicted by the model. The varying parameter in robustness analysis is the logFC threshold to call a gene differentially expressed, as this parameter is regularly set by the user. Tested logFC values were set according to the overall distribution of logFC values between all conditions in the experiment. In particular, quantiles 25, 50 and 75 of the logFC distribution were selected. In addition, logFC equal to 1 was also tested and consequently four different logFC thresholds were evaluated in the robustness analysis. After performing both analyses, highly confidence reaction states were determined consisting of those reactions that pass both the sensitivity and robustness analyses. A reaction has a highly confident or unambiguous state if i) a change of its state reduces the result of the OF, and ii) its state does not change when using different logFC thresholds.

#### 3.4.2 Evaluation of metabolite prediction accuracy

Since the MAMBA model can be used to predict metabolite levels, we used metabolite prediction accuracy to evaluate MAMBA and compare it with MADE, a related approach that incorporates gene expression but no metabolomics data into the GEM. We evaluated the impact of including metabolomic data into the model prediction error using a leave-X-out strategy. Basically, we calculated the error of the MAMBA model in predicting measured metabolites as an increasing number of metabolites were incorporated in the model, i.e., we first fit a model including measurements for one metabolite and compute metabolite prediction error for the remaining metabolite dataset, next we fit the model including measurements for two metabolites and calculate again the error estimates. This was repeated increasing by one the number of included metabolites to reach the complete MAMBA model that includes all available metabolite data and the whole process is repeated one thousand times. The Root Mean Square Error of Prediction (RMSEP) [25] across all predicted metabolites at each leave-X-out iteration was used as error metric.

## 4 Results

### 4.1 MAMBA model

The MAMBA model unlocks the utilization of time-course non-quantitative or relative metabolomics data from different analytic platforms such as MS or NMR as well as widely adopted multi-omics approaches that generate sequencing within the CBM framework for the study of metabolic networks. A number of approaches address partially the integration of these omics into a CBM framework. In this sense D-MFA enables the integration of absolute metabolomics concentration from time-course experiments into a metabolic models, however this approach is limited to quantitative measurements and restricted to networks with low degrees of freedom imposing a strong limitation on the size of the analyzed network. Other methods like MADE are limited to the integration of transcriptomic data from time-course experiments into GEMs. MAMBA addresses this limitation by using relative values whereby the semi-quantitative omics data are modeled as transitions (comparison between two conditions) [7]. Hence, the MAMBA framework uses differential expression/quantification data to characterize the metabolic network across *N* conditions enabling the dynamic modeling of the system. Compared to a basic FBA, MAMBA includes two set of constraints: gene/protein associated constraints and metabolomics associated constraints. This reduces the solution space of the model and the characterization of the metabolic network is more robust and accurate. A basic FBA model is defined by (Figure 1): the stoichiometric matrix, **S**, that indicates the association metabolites (rows) with reactions (columns); vectors **lb** and **ub** that contain the minimal and maximal fluxes allowed for reactions, respectively; vector **b** is the right side of the mass balance equations and is equal to 0 at all its elements to meet the zero mass balance condition; and vector **c** contains the reaction coefficients for the optimization function. The output is a vector **x** with the same size as **c**, i.e., columns of **S**. For MAMBA, gene/protein (via S-GPRs) and metabolomic (via Sink reactions) associated constraints are included alongside **S**, resulting into a new larger matrix, **A**. Figure 3.c shows the design of **A** matrix with a toy metabolic network and considering only two conditions.

### 4.2 Application of MAMBA to a yeast heat-shock dataset improves metabolic prediction accuracy

The time-series multi-omic yeast data (see Section 2) was used to test MAMBA. This is a multifactorial experimental design with two factors: strain (2 levels: WT and *mip*6Δ) and temperature-time (3 levels: baseline or 30 ° C 20 minutes, 39 ° C 20 minutes and 39 ° C 120 minutes) where three omic modalities (RNA-seq, H4K12ac ChIP-seq and metabolomics) were measured. A separate MAMBA model was obtained for each strain, each of them including the three time points of the heat-shock treatment. Consequently, each MAMBA model contains three conditions and two transitions, that is: i) First transition = 30 ° C 20 min. → 39 ° C 20 min., and ii) Second transition = 39 ° C 20 min. → 39 ° C 120 min.. The two MAMBA outputs were then compared to evaluate the differences between strains regarding heat-shock adaptation. Table 1 shows the number of genes, H4K12ac peaks and metabolites measured in our experiment and the number of differentially expressed features that were used to feed the MAMBA models.

**Table 1:**
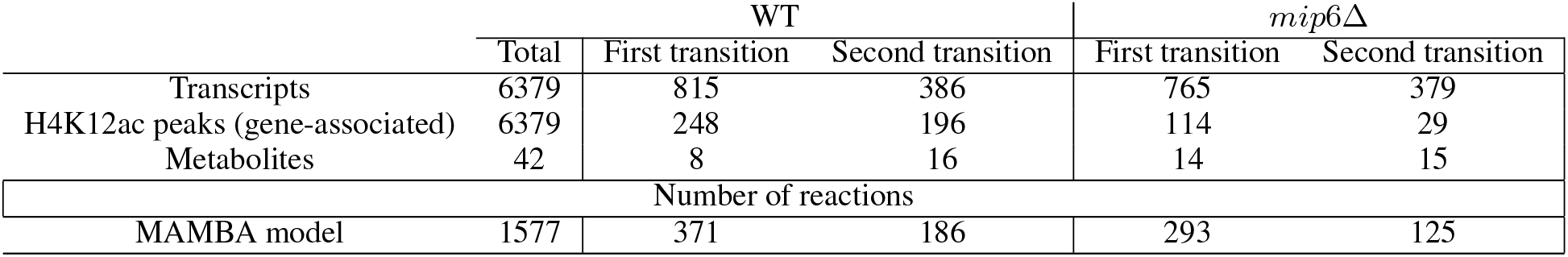
Number of genes, H4K12ac peaks and metabolites measured in our experiment and the number of differentially expressed features (see Methods for details) that were used to feed the MAMBA models.

|  | WT |  |  | *mip6Δ* |  |
| --- | --- | --- | --- | --- | --- |
|  | Total | First transition | Second transition | First transition | Second transition |
| Transcripts | 6379 | 815 | 386 | 765 | 379 |
| H4K12ac peaks (gene-associated) | 6379 | 248 | 196 | 114 | 29 |
| Metabolites | 42 | 8 | 16 | 14 | 15 |
| Number of reactions |  |  |  |  |  |
| MAMBA model | 1577 | 371 | 186 | 293 | 125 |

We used MAMBA results with the WT strain to validate the consistency of the novel approach. First, predicted metabolite ratios were compared with experimental measurements and prediction error was calculated as Root Mean Square Error of Prediction, RMSEP (see section 3.4.2 for details). In addition, MAMBA was compared with MADE algorithm, a similar method that only integrates transcriptomics data [7]. MAMBA and MADE performances were contrasted to evaluate whether the inclusion of a new layer of omic information improves the prediction accuracy of metabolites. We found that MAMBA outperformed MADE in terms of metabolite ratios prediction accuracy with lower RMSEP values in both transitions (Figure 4.a). Moreover, the prediction error decreased when each new metabolite data was incorporated into the model (Figure 4.b). These results demonstrate that the inclusion of metabolite measurements into the GEM boosts network characterization. We also observed that metabolites with other measured metabolites close in the metabolic network (e.g. amino acids), had better predictions than isolated metabolites (Figure 4.c), which adds to the consistency of the MAMBA model when incorporating metabolic information.

**Figure 4:**
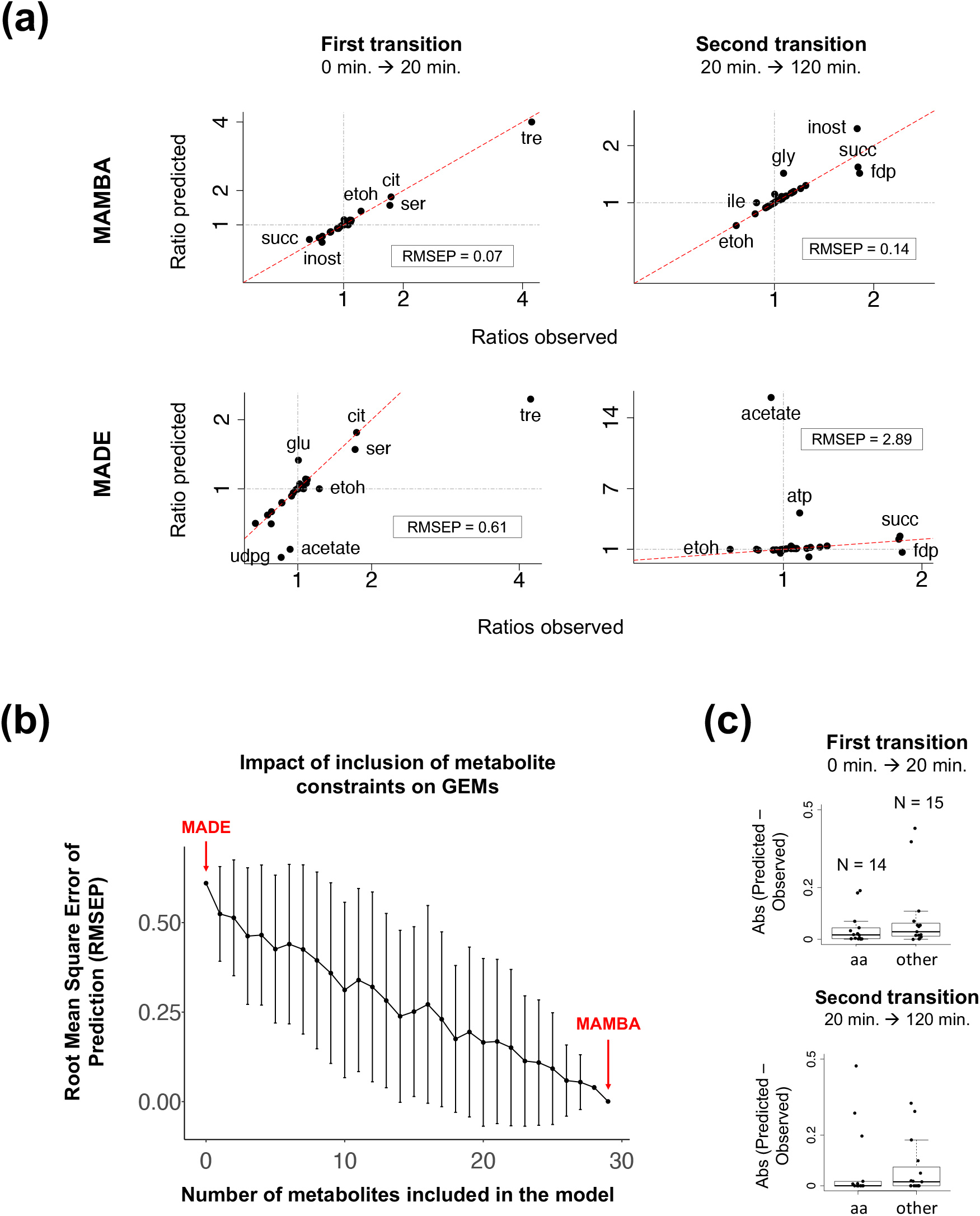
MAMBA validation of prediction accuracy. a) Metabolite prediction accuracy. Predicted vs observed metabolite ratios are shown for both transitions (comparisons) in the WT model. MAMBA is compared to MADE which only uses transcriptomic data. b) The effect of the number of metabolites included in the model on the prediction error (RMSEP) only for the first transition. Mean (dots) sd ±(error bars) are represented. c) Prediction error of MAMBA by metabolite type separating between amino acids (aa) and other metabolites.

MAMBA results also recapitulated the known biology. We delved into the Trehalose pathway, which is a well-known process affected by heat stress in yeast cells [26]. MAMBA predicted the activation of Trehalose production pathway after heat-shock and its maintenance at 120 minutes (Figure 5.a). However, trehalose metabolism genes were down-regulated after 120 minutes compared to 20 minutes of heat stress (Figure 5.b) which was not readily consistent with high trehalose production at 120 minutes. Nonetheless, our metabolomic data show trehalose rises steadily during heat stress (Figure 5.c1). Indeed, MAMBA predicted Trehalose over-production (Figure 5.d) regardless of gene expression data. Again, this is consistent with the inclusion of metabolomics data improving network characterization as only gene expression is not always completely representative of metabolic changes.

**Figure 5:**
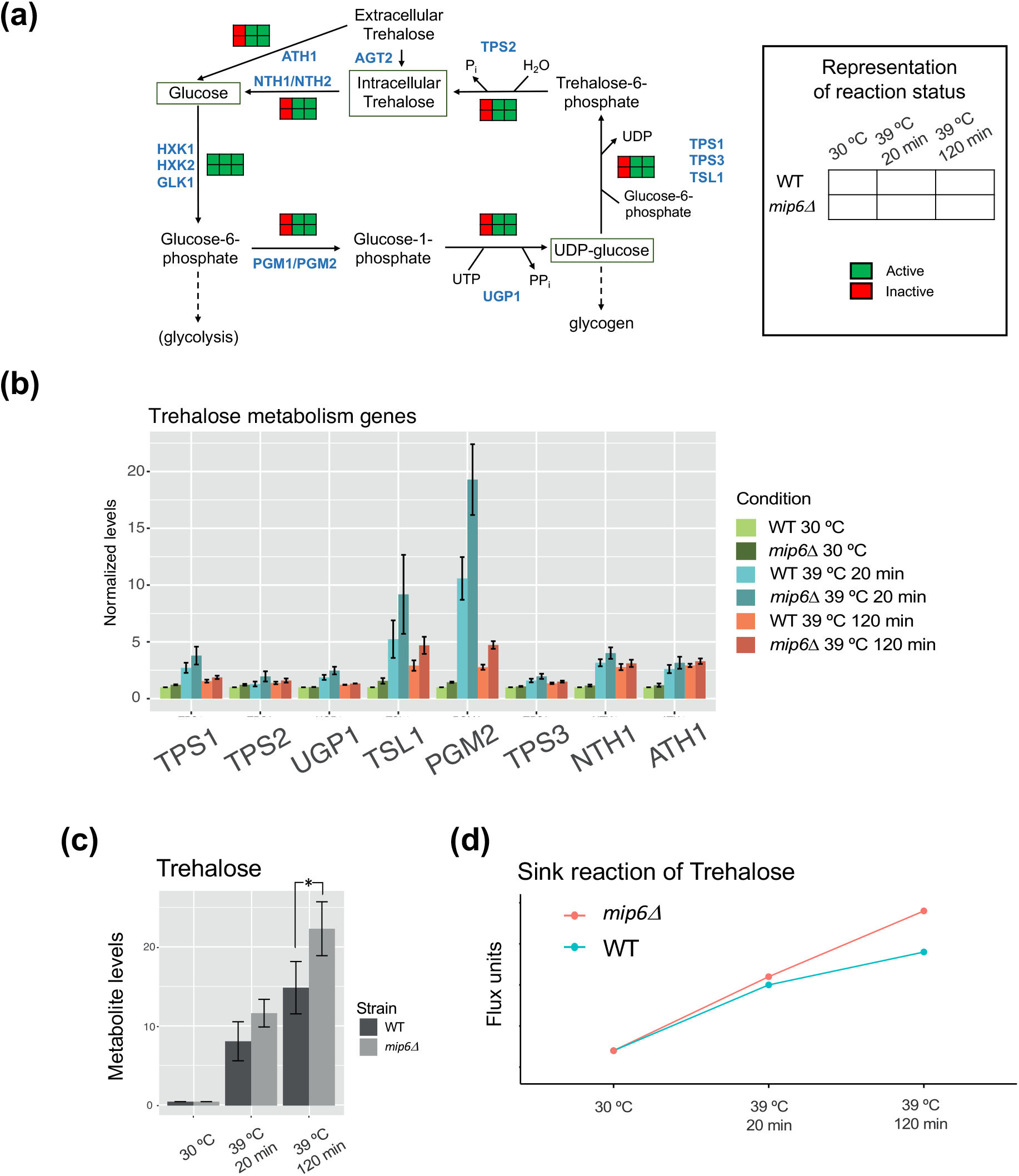
MAMBA validation of biological consistency. a) Trehalose pathway: Predicted reaction status. Trehalose pathway activity is predicted to increase at 20 min and to remain active allowing trehalose accumulation. b) Expression of genes involved in trehalose pathway. c) Trehalose quantification from NMR metabolomic data. d) MAMBA prediction of Trehalose quantification.

### 4.3 Deciphering differential behavior between strains

We next used MAMBA to study metabolic differences between strains. Reaction states (active or inactive) were compared, and we found 211 reactions that had a different state between strains at one time point at least (list of reactions at Supplementary Table 1). To understand the biological meaning of these differences, we performed a pathway-level analysis where yeast pathways from KEGG database were used. Given a metabolic pathway *P* containing *n* reactions, we defined a pathway enrichment score (PES) as the percentage of reactions of *P* contained in the list of differential reactions identified by MAMBA (*Q*):

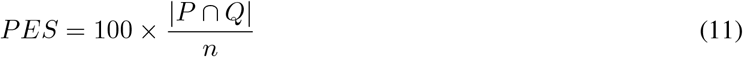

In order to set a relevance threshold, we compared each pathway PES to the PES in the general KEGG pathway, “Metabolic Pathways” -that contains all metabolic reactions-, which was 17%. Thus, those pathways with a PES higher than 17% were selected as relevant pathways to describe strain differences. This resulted into a list of 25 pathways showed in Figure 6.a. We observed that this relevant pathways recapitulate main biological changes across conditions using Gene Set Variation Analysis (GSVA) method [27], that computes a sample-wise score for each pathway that summarizes the expression of the genes contained in it. An unsupervised clustering performed on the GSVA scores Figure 6.a showed that samples (columns) were separated by both experimental conditions (strain and time), indicating that the list of relevant pathways summarized the main biological signal across conditions.

**Figure 6:**
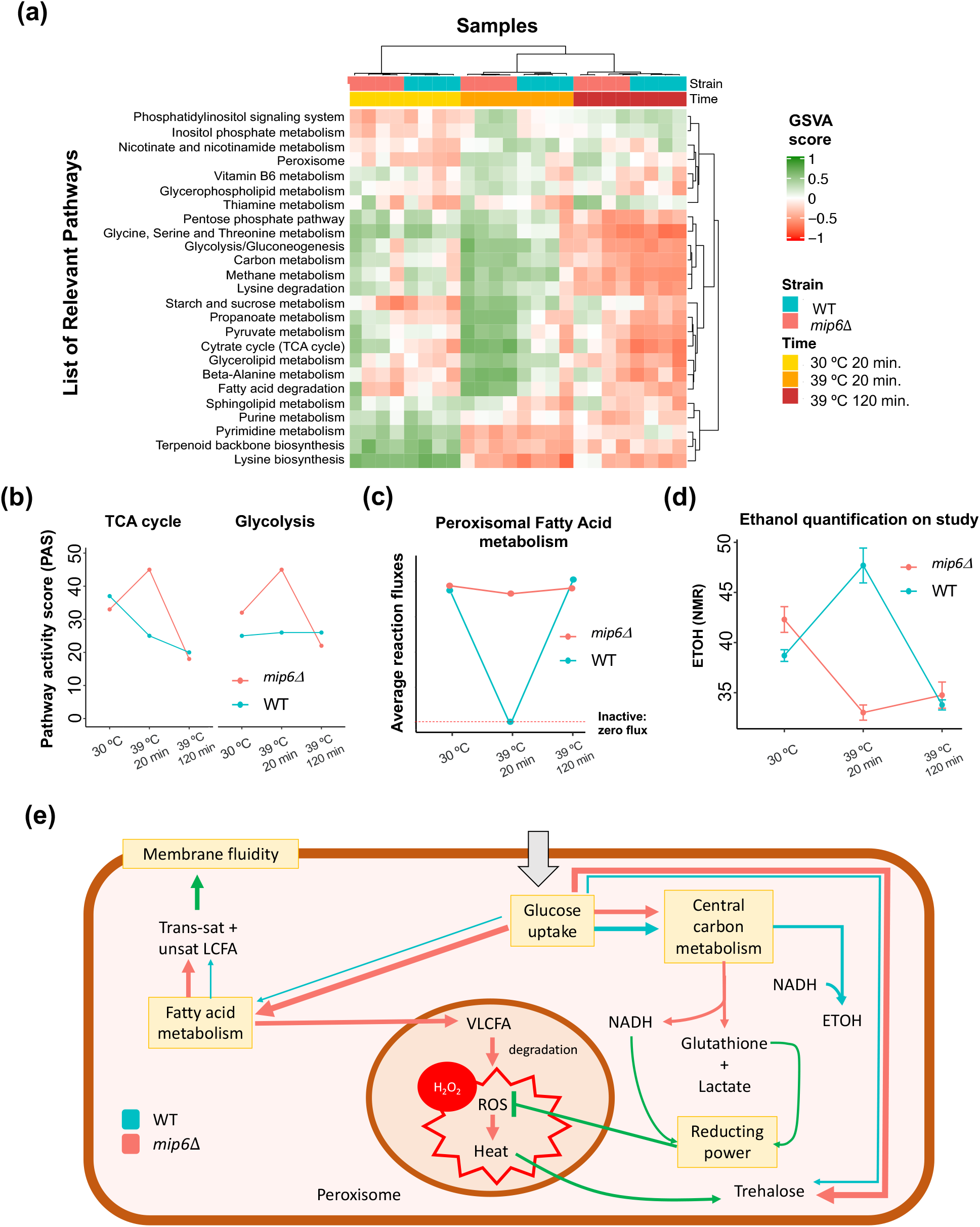
Model of differential adaptation to heat-shock between strains revealed by MAMBA analysis. a) Relevant pathways heatmap. Those pathways containing reactions with differential activity state profile between strains were considered. Samples are fully separated according to experimental conditions. b) Pathway activity score of Glycolisis and TCA cycle, calculated as the percentage of active reactions. c) MAMBA predicted fluxes through peroxisomal fatty acid metabolism reactions. MAMBA predicts a shutdown of this pathway in WT while it remains active in the mutant. d) Ethanol quantification from NMR metabolomics data. e) Explanatory model of metabolic differences found between strains. From the carbon source (glucose), the WT produces ethanol, trehalose and LCFAs (Long-chain Fatty acids), that contributes to membrane stabilization. On the contrary, *mip*6Δ mutant produces lactate + glutathione instead of ethanol, a higher amount of trehalose and LCFAs, which are metabolized in the peroxisome to produce H_2_O_2_ resulting in oxidative and heat stress for the mutant cells.

To better represent the predicted activity for these relevant pathways, we computed the pathway activity scores (PAS) that is the percentage of active reactions (binary reaction status equal to 1):

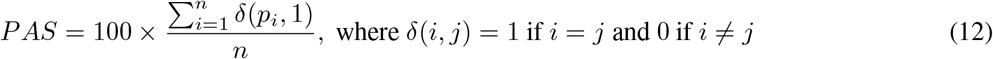

Figure 6.b represents PAS of two key processes within core carbon metabolism, Glycolysis and TCA cycle. Activity scores for the rest of relevant pathways can be found in Supplementary Figure 1. We observed that *mip*6Δ mutant over-activates Glycolysis and TCA cycle while the WT decreases the activity of TCA cycle and maintain the status of Glycolysis. Similar to TCA, Pyruvate metabolism is down-regulated in WT but over-activated in the mutant during heat stress. In fact, global “Carbon metabolism” pathway presented the same pattern, and “Peroxisome metabolism” and “Pentose phosphate pathway” also showed similar profile. “Fatty Acid (FA) degradation” profile was also different between strains. WT maintains “FA degradation” over time while the mutant showed a huge increase of FA degradation activity. Interestingly, “FA degradation” and “Peroxisome metabolism” shared most of the mapped differential reactions that were involve in Peroxisomal FA degradation (PFAD). Analyzing the average flux through reaction involved in PFAD we observed a consistent pattern where the average flux decreases in the WT and stay flat in the mutant (Figure 6.c). Lastly, activation scores of “Purine” and “Pyrimidine” metabolism pathways slightly decrease in the WT and remains flat in the mutant.

The disconnection of TCA cycle and Glycolysis in the WT can be explained by the increase in Ethanol (ETOH) production after heat-shock (Figure 6.d). MAMBA predicts this differential behavior as the modeled ethanol ratios for both transitions are 1.4 (time 20 vs time 0) and 0.6 (time 120 vs time 20) for the WT (Figure 4.a).

Based on the list of differential reactions we then evaluated which reactants and products were predominant. Table 2 shows that NADH/NADPH were enriched as products of differential reactions, and therefore NAD/NADP were enriched as reactants. Moreover, most of the reactions that have either NADH or NADPH as products (12/15) were down-regulated in the WT after heat-shock but not in the mutant. This result suggested an imbalance in the generation of reducing power between strains towards *mip*6Δ mutant.

**Table 2:** Number of reactions that have NADH or NADPH as products (left) and NAD or NADP as reactants (right) separated by whether they are in the list of 211 differential reactions between strains. Significant enrichment assessed by Fisher’s exact test [28]

|  | NADH/NADPH as products |  | NAD/NADP as reactants |  |
| --- | --- | --- | --- | --- |
|  | Yes | No | Yes | No |
| Differential Reactions | 15 | 198 | 15 | 198 |
| Rest | 31 | 1333 | 35 | 1329 |
| Fisher’s exact test p-val: | 0.0006 |  | 0.002 |  |

An explanatory model of the main metabolic differences between strains found by MAMBA is represented in Figure 6.e. Starting from the carbon source (glucose), the WT produces ethanol, trehalose and LCFAs (Long-chain Fatty acids), that contributes to membrane stabilization [29]. On the contrary, *mip*6Δ mutant produces lactate + glutathione instead of ethanol, a higher amount of trehalose and LCFAs, which are metabolized in the peroxisome to produce H_2_O_2_ resulting in oxidative and heat stress for the mutant cells.

### 4.4 Evaluating the effect of mip6 affinity on metabolic changes

We seized PAR-CLIP data, a technology that profiles RNA-protein interactions, to assess to which extent metabolic differences between strains might be caused by direct interaction with mip6. Mip6 is involved in RNA export from the nucleus and in stabilizing mRNAs in the cytosol through direct protein-RNA interactions [20]. PAR-CLIP data was available for WT yeast strain on the same heat stress conditions, indicating a total of 6685 mRNAs bound by mip6, with a small fraction of them (488) showing differential mip6 affinity after heat stress [20]. We compared this list of mip6-bound RNAs to the 150 genes involed in the differential reactions identified by MAMBA. We found that this set of genes showed higher mip6 affinity both at normal growth condition (30 ° C) and after heat-shock (39 ° C) (Figure 7.a and b) than genes not involved in MAMBA-detected reactions. However, only 10 genes showed differential mip6 affinity at 39 ° C compared to 30 ° C (Supplementary Table 2).

**Figure 7:**
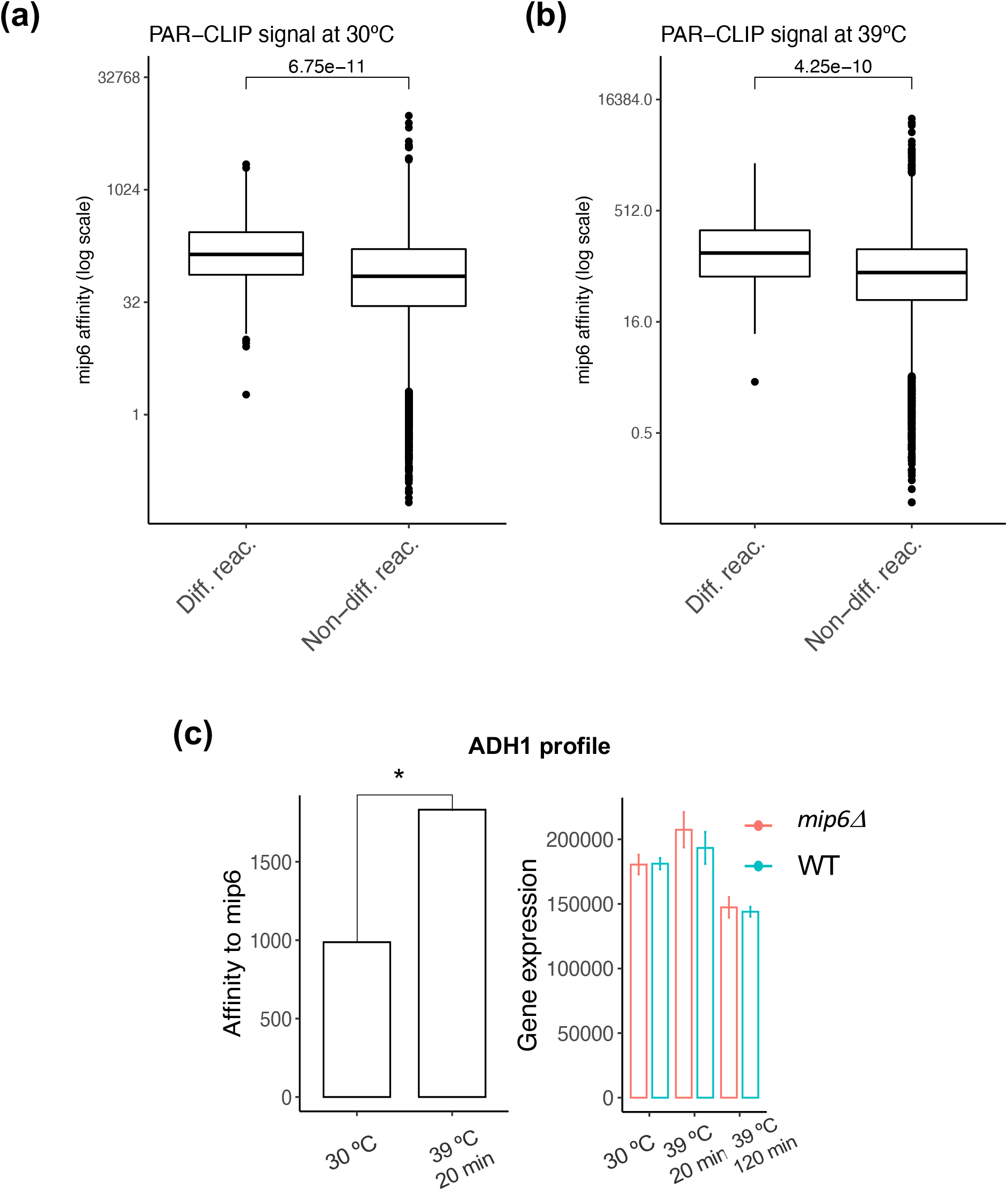
Evaluation of mip6 PAR-CLIP data from Martin-Exposito et al., 2019 [20]. a) Signal at 30 ° C (normal growth condition). b) Signal at 39 ° C (heat stress). Mip6 affinity is compared between genes involved in differential reactions identified by MAMBA and the rest of genes. Wilcoxon-test p-value is shown [30, 31]. c) ADH1 profile. [left panel] PAR-CLIP data reflecting ADH1 affinity to mip6, which increases during heat-shock. [right panel] ADH1 expression with no significant differences between strains.

One of the genes identified by the MAMBA analysis which showed a differential mip6 affinity upon heat-shock was ADH1. This gene, responsible for the reduction of acetaldehyde to ethanol, showed a strong mip6 affinity increase at 39°C (Figure 7.c) what might explain its stabilization in the WT and therefore the increase in ETOH production. Interestingly, ADH1 expression did not change between strains after heat stress which indicates that the lower ETOH production in the mutant is not caused by differences in gene expression (Figure 7.c).

### 4.5 Metabolic control by ChIP-seq signal

In order to understand the contribution of histone modifications to metabolic control, MAMBA was run using ChIP-seq data instead of gene expression to predict reaction fluxes. Firstly, we compared reaction status between RNA-seq and ChIP-seq driven MAMBA models and found a higher consistency for the WT as the status of 59.8% of reactions were equally derived from both input data types, while both outputs for *mip*6Δ mutant showed a consistency of 40.2% (Table 4). MAMBA model using ChIP-seq data also showed a huge difference between WT and *mip*6Δ regarding down-regulated reactions after heat-shock, i.e. the mutant presented a lower number of reactions that became inactive after heat-shock according to ChIP-seq data.

We also evaluated ChIP-seq data of those genes contained in the list of relevant pathways (Figure 6.e), and we found statistically significant differences between strains regarding Peroxisomal FA metabolism (Figure 8.a). WT ChIP-seq signal of this set of genes indicated a decrease in histone acetylation signal, that was consistent with the down-regulation of gene expression and the corresponding inactivation of these metabolic reactions and of the flux through the pathway. On the contrary, ChIP-seq data in *mip*6Δ mutant did not change. Figure 8.a and Table 4 show that *mip*6Δ mutant has a lower number of de-acetylation changes compared to the WT. This difference between strains was not due to gene expression differences of yeast HDACs (Histone Deacetylases) as they were similarly expressed in both strains (Figure 8.b).

**Figure 8:**
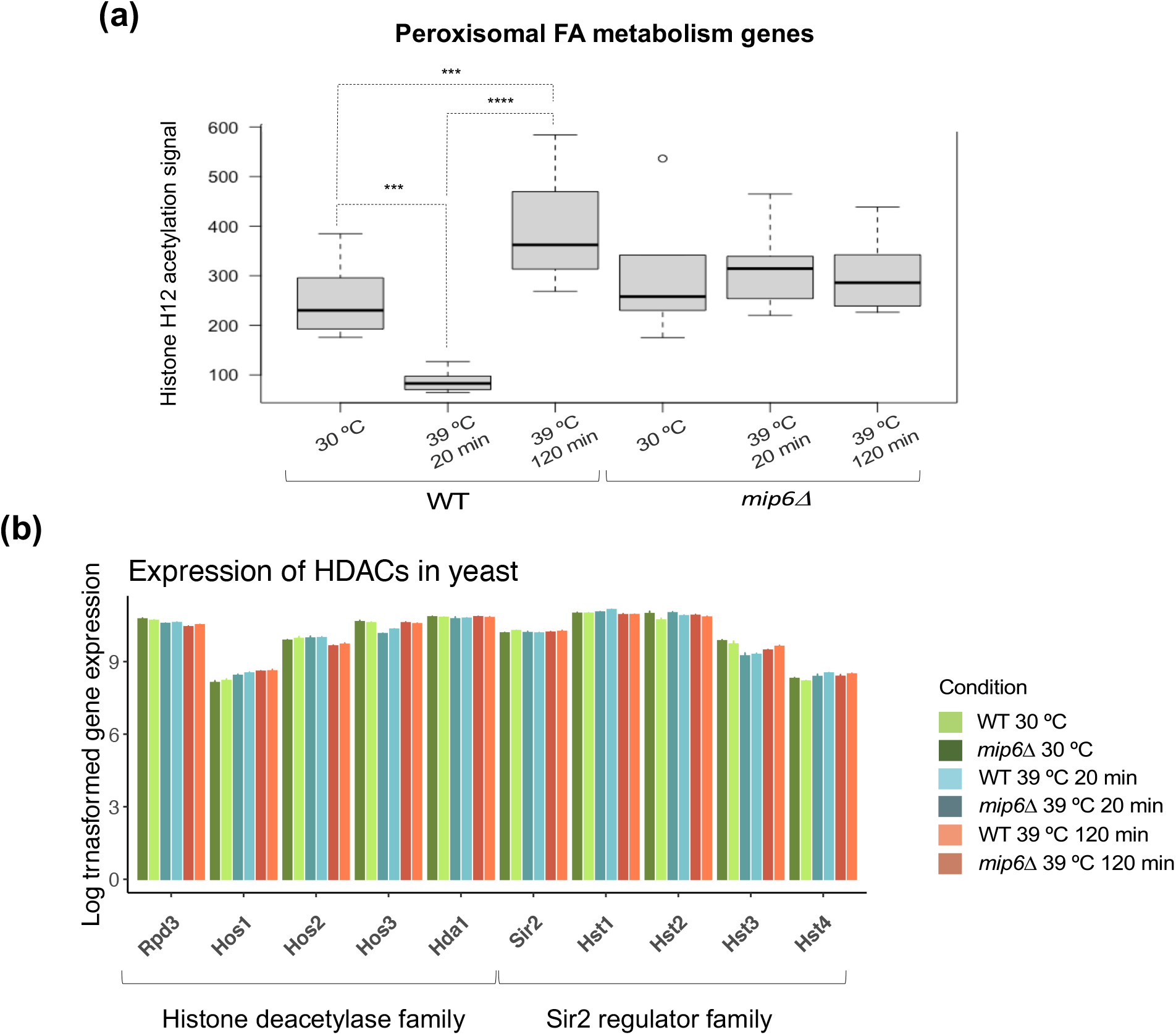
Peroxisomal Fatty acid metabolism control by ChIP-seq signal. a) ChIP-seq signal of genes involved in peroxisomal FA metabolism. WT underwent de-acetylation of those genes while no changes were present in the mutant. Statistical significance determined by Wilcoxon-Mann Whitney test [30, 31] (***: p-value < .001; ****: p-value < .0001). b) Expression of the HDACs in *S. cerevisiae*. No statistically significant differences between strains were found.

## 5 Discussion

In this work we have introduced MAMBA (Metabolic Adjustment via Multiomic Block Aggregation), a constraint-based genome-scale metabolic reconstruction algorithm that allows the integration of gene-associated omic data and semi-quantitative metabolomics. Compared to previous approaches, MAMBA has a better metabolite prediction accuracy, which means a more accurate metabolic network characterization. Additionally, MAMBA simultaneously models multiple conditions and can therefore be applied to the analysis of time-course data, providing a modelling framework for dynamic processes. In this work, we applied MAMBA to study metabolic regulation in two yeast strains (WT and the *mip*6Δ mutant) after a heat-shock treatment.

The output of MAMBA recapitulates the known yeast behavior under heat stress condition. Specifically, trehalose production is one of the most important heat stress response mechanisms and behaves as a heat protector in yeast [26].

Trehalose production increases under heat stress condition [32] and has been shown to be a powerful stabilizer of proteins and membranes [33]. Only considering transcriptomics data, our data showed that trehalose production reaches its maximum at 20 minutes after the heat-shock and then it starts decreasing to show at 120 minutes similar values to the initial ones. However, trehalose is accumulated in cells according to metabolomics data and MAMBA was able to reveal this behavior. Therefore, MAMBA is able to leverage metabolomics data to improve overall metabolite prediction accuracy beyond gene expression changes. Importantly, MAMBA is also useful for driving novel biological findings. We focused on comparing both strains in terms of heat stress adaptation. Using MAMBA, we have identified the underlying mechanism that explains the differential dynamics between strains both for transcriptomic and metabolomic changes. Overall, our results suggest that the mutated strain has a lower capacity to adapt to heat stress compared to the WT. We found that Carbon metabolism (Glycolysis, Pyruvate metabolism and TCA cycle among others) was differentially regulated between strains, and in consequence, both strains adapt differently to the heat-shock treatment. Results indicate a disconnection between TCA cycle and Glycolysis in WT while the mutant requires more flux through these pathways that could be related to energetic needs. This example also demonstrates the advantages of model-driven approaches over traditional gene expression analysis. Following a standard differential expression and functional enrichment analysis, TCA cycle was not identified as being differentially regulated between the two strains (Supplementary Table 3), possibly because only 3 out of 31 genes annotated in TCA cycle were differentially expressed, which are too few to support a significant enrichment result. However, reactions codified by those three genes are critical in the TCA cycle: Pyruvate Dehydrogenase (pyruvate to ac-CoA), Citrate Synthase (Oxoglutarate to Citrate) and Aconitase (Citrate to Isocitrate) and this critical contribution of these reactions to the pathway activity was captured by the metabolic modelling.

**Table 3:** Number of reactions that change over time in both strains according RNA-seq and ChIP-data.

| Reaction profile |  | RNA-seq model |  | ChIP-seq model |  | Overlap |  |
| --- | --- | --- | --- | --- | --- | --- | --- |
| First transition | Second transition | WT | *mip6*Δ | WT | *mip6*Δ | WT | *mip6*Δ |
| Constant | Decrease | 96 | 57 | 75 | 23 | 64 | 14 |
| Constant | Increase | 90 | 68 | 77 | 59 | 53 | 32 |
| Increase | Constant | 68 | 118 | 57 | 101 | 42 | 80 |
| Increase | Decrease | 69 | 99 | 12 | 48 | 8 | 37 |
| Decrease | Constant | 93 | 54 | 86 | 19 | 75 | 5 |
| Decrease | Increase | 141 | 22 | 103 | 0 | 91 | 0 |
| Total |  | 557 | 418 | 410 | 250 | 333 | 168 |

Moreover, the analysis at reaction level returned important differences between strains regarding heat stress adaptation. MAMBA indicated that *mip*6Δ mutants fail to shut the peroxisomal fatty acid metabolism down which causes an increased oxidative and heat stress. One subproduct of the peroxisomal FA degradation is Hydrogen Peroxide (*H*_2_*O*_2_) which generates heat when it is metabolized in the cell. This may explain why the *mip*66Δ continues generating reducing power after the heat-shock (Table 2) as well as why the mutant also produces more Trehalose (heat protective effect) during heat-shock. Additionally, the MAMBA model based on H4K12ac data indicated that the inactivation of peroxisomal FA metabolism in the WT was controlled epigenetically, which is consistent with previous studies showing that the response to heat stress is associated to changes in chromatin acetylation [19]. Globally, the ChIP-seq MAMBA model indicated that the mutant is less efficient in chromatin de-acetylation not only for peroxisomal FA metabolism genes but in general for the whole genome-scale model. Importantly, this observation in the mutant is not caused by a lower expression of HDACs as they are not under-expressed in the mutant. This opens up the question of the existence of a potential histone de-acetylation regulatory mechanism beyond gene expression of effector proteins, that might be linked to mip6.

Ethanol production during heat-shock was also different between strains. Ethanol production increased in WT at time 20min after heat-shock while it decreased in the mutant. Interestingly, *mip*6Δ increases lactate production while WT does not [19]. In the metabolic model used, lactate can only be produced by a reaction that also produces glutathione, which is a well known source of cellular reducing power. Additionally, ethanol production in yeast consumes reducing power in the form of NADH. Therefore, the difference in the ethanol/lactate production between strains could also be linked to peroxisomal FA metabolism. Interestingly, PAR-CLIP data on the same WT strain revealed a direct interaction between ADH1 and mip6 that could be related to the known role of mip6 as an mRNA export factor and mRNA stabilizer in the cytoplasm and an alternative explanation of ethanol profile in the mutant compared to the WT. Proline is also able to confer heat protection to yeast cells [34], however its production decreases during heat-shock in wild type yeast [32]. Proline production increased in the *mip*6Δ under heat stress strengthening the hypothesis of a higher-stressed mutants compared to WT cells [19].

All together, the MAMBA model of the *mip*6Δ mutant heat-response suggests that the metabolism of the mutant is less flexible to environmental challenges which may be related to the roles of mip6 in the cell: i) interaction with HDACs and ii) mRNA export and stabilizer. As a consequence, mip6 knockdown results into a hyper-stressed state compared to WT and therefore the key yeast thermal protectors, trehalose and proline, are overproduced in the mutant.

In conclusion, we have demonstrated MAMBA is a powerful methodology for constructing robust metabolic networks. MAMBA outperforms other GEMs in terms of metabolite prediction accuracy and is useful to reveal patterns of metabolic control from the combination of matching transcriptomic and metabolomic data. Moreover, MAMBA allows the dynamic modeling of the system and therefore is specially powerful to analyze time-series data and compare over-time profiles among different conditions.

