## Supplementary Material for "MAMBA: a model-driven, constraint-based multiomic integration method"

### 6 Supplementary material

#### 6.1 Supplementary Figure 1

Pathway Activation Score (PAS) for the list of relevant pathways.

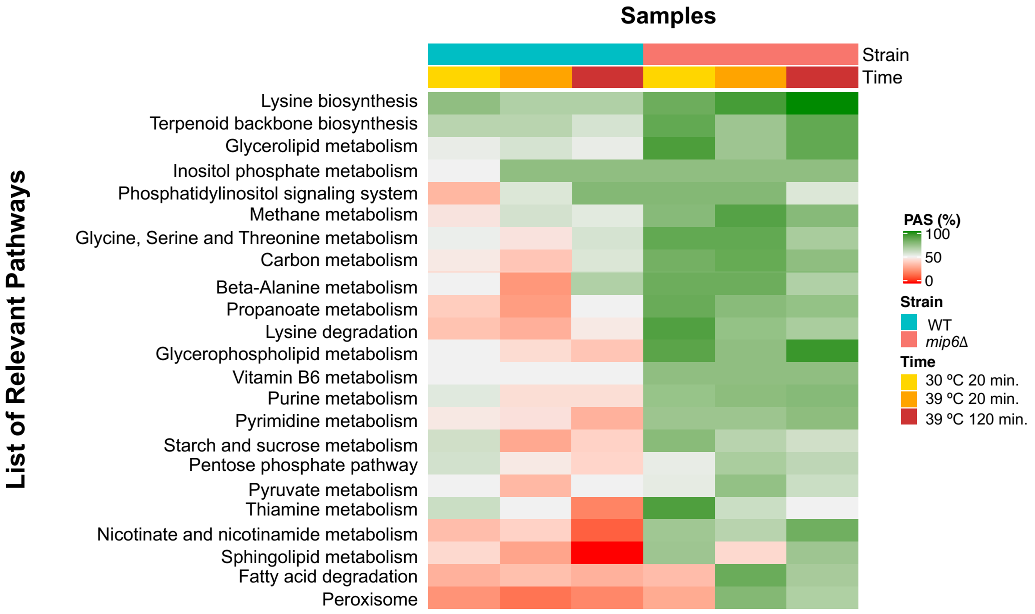

##### 6.2 Supplementary Table 1

List of differential reactions identified by MAMBA and their activation state.

| Reaction data | | | | Wild Type | | | $mip6\Delta$ | | |
|---|---|---|---|---|---|---|---|---|---|
| Rxn | Reactants | Products | Pathways | t0 | t20 | t120 | t0 | t20 | t120 |
| 2MBALDt | M-2mbald-c | M-2mbald-e | NA | 1 | -1 | -1 | -1 | 1 | 1 |
| 2MBTOHtm | M-2mbtoh-c | M-2mbtoh-m | NA | -1 | 1 | -1 | 1 | 1 | 1 |
| AASAD1 | M-L2aadp-c M-atp-c M-h-c M-nadph-c | M-L2aadp6sa-c M-TM-atp-c M-amp-c M-nadp-c M-ppi-c | sce00300-Lysine-biosynthesis sce00770-Pantothenate-and-CoA-biosynthesis sce01100-Metabolic-pathways sce01110-Biosynthesis-of-secondary-metabolites sce01230-Biosynthesis-of-amino-acids | -1 | 1 | -1 | 1 | 0 | 1 |
| AASAD2 | M-L2aadp-c M-atp-c M-h-c M-nadh-c | M-L2aadp6sa-c M-TM-atp-c M-TM-nad-c M-amp-c M-nad-c M-ppi-c | sce00300-Lysine-biosynthesis sce00770-Pantothenate-and-CoA-biosynthesis sce01100-Metabolic-pathways sce01110-Biosynthesis-of-secondary-metabolites sce01230-Biosynthesis-of-amino-acids | -1 | -1 | 1 | -1 | 1 | 1 |
| ACACT1m | M-accoa-m | M-aacoa-m M-coa-m | sce00071-Fatty-acid-degradation sce00072-Synthesis-and-degradation-of-ketone-bodies sce00280-Valine,-leucine-and-isoleucine-degradation sce00310-Lysine-degradation sce00380-Tryptophan-metabolism sce00620-Pyruvate-metabolism sce00630-Glyoxylate-and-dicarboxylate-metabolism sce00640-Propanoate-metabolism sce00650-Butanoate-metabolism sce00900-Terpenoid-backbone-biosynthesis sce01100-Metabolic-pathways sce01110-Biosynthesis-of-secondary-metabolites sce01200-Carbon-metabolism sce01212-Fatty-acid-metabolism | 1 | 1 | -1 | 1 | 0 | 1 |
| ACHLE1 | M-h2o-c M-iamac-c | M-TM-ac-c M-ac-c M-h-c M-iamoh-c | NA | 1 | 1 | -1 | 1 | 0 | 1 |
| ACHLE2 | M-h2o-c M-ibutac-c | M-TM-ac-c M-ac-c M-h-c M-ibutoh-c | NA | -1 | 1 | -1 | 1 | 1 | 1 |
| ACHLE3 | M-aces-c M-h2o-c | M-TM-ac-c M-TM-etoh-c M-ac-c M-etoh-c M-h-c | NA | -1 | 1 | -1 | 1 | 1 | 1 |

| Reaction data | | | | Wild Type | | | $mip6\Delta$ | | |
|---|---|---|---|---|---|---|---|---|---|
| Rxn | Reactants | Products | Pathways | t0 | t20 | t120 | t0 | t20 | t120 |
| ADA | M-adn-c M-h2o-c M-h-c | M-ins-c M-nh4-c | sce00230-Purine-metabolism sce01100-Metabolic-pathways | -1 | 1 | -1 | -1 | 1 | 1 |
| ADK1m | M-amp-m M-atp-m | M-adp-m | sce00230-Purine-metabolism sce00730-Thiamine-metabolism sce01100-Metabolic-pathways sce01110-Biosynthesis-of-secondary-metabolites | 1 | 1 | -1 | 1 | 1 | 1 |
| ADK4m | M-amp-m M-itp-m | M-adp-m M-idp-m | sce00230-Purine-metabolism sce00730-Thiamine-metabolism sce01100-Metabolic-pathways sce01110-Biosynthesis-of-secondary-metabolites | -1 | -1 | 1 | 1 | 1 | 1 |
| ADNCYC | M-atp-c | M-TM-atp-c M-camp-c M-ppi-c | sce00230-Purine-metabolism sce01100-Metabolic-pathways sce04113-Meiosis sce04213-Longevity-regulating-pathway | 1 | -1 | -1 | 1 | 1 | 1 |
| ADNK1 | M-adn-c M-atp-c | M-TM-atp-c M-adp-c M-amp-c M-h-c | sce00230-Purine-metabolism sce01100-Metabolic-pathways | -1 | 1 | 1 | 1 | 1 | 1 |
| ADNUC | M-adn-c M-h2o-c | M-ade-c M-rib–D-c | sce00240-Pyrimidine-metabolism sce00760-Nicotinate-and-nicotinamide-metabolism sce01100-Metabolic-pathways | 1 | 1 | -1 | 1 | 1 | 1 |
| AGTi | M-ala–L-c M-glx-c | M-TM-ala–L-c M-TM-gly-c M-gly-c M-pyr-c | sce00250-Alanine,-aspartate-and-glutamate-metabolism sce00260-Glycine,-serine-and-threonine-metabolism sce00630-Glyoxylate-and-dicarboxylate-metabolism sce00680-Methane-metabolism sce01100-Metabolic-pathways sce01110-Biosynthesis-of-secondary-metabolites sce01200-Carbon-metabolism sce04146-Peroxisome | -1 | 1 | 1 | 1 | 1 | 1 |
| ALCD2x-copy1 | M-etoh-c M-nad-c | M-TM-etoh-c M-TM-nad-c M-acald-c M-h-c M-nadh-c | sce00010-Glycolysis-/-Gluconeogenesis sce00071-Fatty-acid-degradation sce00350-Tyrosine-metabolism sce01100-Metabolic-pathways sce01110-Biosynthesis-of-secondary-metabolites | -1 | 1 | 1 | 1 | 1 | 1 |

| Reaction data | | | | Wild Type | | | $mip6\Delta$ | | |
|---|---|---|---|---|---|---|---|---|---|
| Rxn | Reactants | Products | Pathways | t0 | t20 | t120 | t0 | t20 | t120 |
| ALDD2ym | M-acald-m M-h2o-m M-nadp-m | M-ac-m M-h-m M-nadph-m | sce00010-Glycolysis-/-Gluconeogenesis sce00071-Fatty-acid-degradation sce00280-Valine,-leucine-and-isoleucine-degradation sce00310-Lysine-degradation sce00330-Arginine-and-proline-metabolism sce00340-Histidine-metabolism sce00380-Tryptophan-metabolism sce00410-beta-Alanine-metabolism sce00561-Glycerolipid-metabolism sce00620-Pyruvate-metabolism sce00770-Pantothenate-and-CoA-biosynthesis sce01100-Metabolic-pathways sce01110-Biosynthesis-of-secondary-metabolites | -1 | 1 | -1 | 1 | 1 | 1 |
| AMPN | M-amp-c M-h2o-c | M-ade-c M-r5p-c | sce00230-Purine-metabolism sce01100-Metabolic-pathways sce01110-Biosynthesis-of-secondary-metabolites | 1 | -1 | -1 | 1 | 1 | 1 |
| ASNN | M-asn–L-c M-h2o-c | M-TM-asn–L-c M-TM-asp–L-c M-asp–L-c M-nh4-c | sce00250-Alanine,-aspartate-and-glutamate-metabolism sce00460-Cyanoamino-acid-metabolism sce01100-Metabolic-pathways sce01110-Biosynthesis-of-secondary-metabolites | 1 | -1 | -1 | 1 | 1 | 1 |
| ASPt2n | M-asp–L-c M-h-c | M-TM-asp–L-c M-asp–L-n M-h-n | NA | -1 | 1 | -1 | 1 | 1 | 1 |
| ASPt2r | M-asp–L-e M-h-e | M-TM-asp–L-c M-asp–L-c M-h-c | NA | 1 | -1 | 1 | 1 | 1 | 1 |
| ATPH1 | M-atp-c M-h2o-c | M-TM-atp-c M-amp-c M-h-c M-pi-c | sce00230-Purine-metabolism sce00240-Pyrimidine-metabolism sce01100-Metabolic-pathways | 1 | -1 | 1 | 1 | 1 | 1 |
| ATPM | M-atp-c M-h2o-c | M-TM-atp-c M-adp-c M-h-c M-pi-c | NA | 1 | -1 | -1 | -1 | -1 | 1 |
| ATPS3m | M-adp-m M-h-c M-pi-m | M-atp-m M-h2o-m M-h-m | sce00190-Oxidative-phosphorylation sce01100-Metabolic-pathways | -1 | 0 | 1 | 1 | 1 | 1 |
| BTDD-RR | M-btd-RR-c M-nad-c | M-TM-nad-c M-actn–R-c M-h-c M-nadh-c | sce00650-Butanoate-metabolism | -1 | -1 | -1 | -1 | 0 | 1 |
| CERASE224er | M-cer2-24-r M-coa-r M-h-r | M-psphings-r M-ttccoa-r | sce00600-Sphingolipid-metabolism sce01100-Metabolic-pathways | 1 | 1 | -1 | 1 | 1 | 1 |

| Reaction data | | | | Wild Type | | | $mip6\Delta$ | | |
|---|---|---|---|---|---|---|---|---|---|
| Rxn | Reactants | Products | Pathways | t0 | t20 | t120 | t0 | t20 | t120 |
| CERS224er | M-psphings-r M-ttccoa-r | M-cer2-24-r M-coa-r M-h-r | sce00600-Sphingolipid-metabolism sce01100-Metabolic-pathways | 1 | 1 | -1 | 1 | 1 | 1 |
| CHLPCTD | M-cholp-c M-ctp-c M-h-c | M-cdpchol-c M-ppi-c | sce00440-Phosphonate-and-phosphinate-metabolism sce00564-Glycerophospholipid-metabolism sce01100-Metabolic-pathways | -1 | -1 | 1 | 1 | 1 | 1 |
| CO2tm | M-co2-c | M-co2-m | NA | -1 | 1 | 1 | 1 | 1 | 1 |
| COAtim | M-coa-c | M-coa-m | NA | 1 | -1 | 1 | 1 | 1 | 1 |
| CTPS2 | M-atp-c M-gln–L-c M-h2o-c M-utp-c | M-TM-atp-c M-TM-gln–L-c M-TM-glu–L-c M-adp-c M-ctp-c M-glu–L-c M-h-c M-pi-c | sce00240-Pyrimidine-metabolism sce01100-Metabolic-pathways | -1 | 1 | 1 | 1 | 1 | 1 |
| CYSS | M-acser-c M-h2s-c | M-TM-ac-c M-ac-c M-cys–L-c M-h-c | sce00270-Cysteine-and-methionine-metabolism sce00920-Sulfur-metabolism sce01100-Metabolic-pathways sce01110-Biosynthesis-of-secondary-metabolites sce01200-Carbon-metabolism sce01230-Biosynthesis-of-amino-acids | 1 | -1 | 1 | 1 | 1 | 1 |
| CYTD | M-cytd-c M-h2o-c M-h-c | M-nh4-c M-uri-c | sce00240-Pyrimidine-metabolism sce01100-Metabolic-pathways | 1 | 1 | -1 | 1 | 1 | 1 |
| CYTK1 | M-atp-c M-cmp-c | M-TM-atp-c M-TM-cmp-c M-adp-c M-cdp-c | NA | -1 | -1 | -1 | 1 | -1 | 1 |
| DHAK | M-atp-c M-dha-c | M-TM-atp-c M-adp-c M-dhap-c M-h-c | sce00051-Fructose-and-mannose-metabolism sce00561-Glycerolipid-metabolism sce00680-Methane-metabolism sce01100-Metabolic-pathways sce01200-Carbon-metabolism | 1 | -1 | 1 | 1 | 1 | 1 |
| DURIPP | M-duri-c M-pi-c | M-2dr1p-c M-ura-c | sce00230-Purine-metabolism sce00240-Pyrimidine-metabolism sce00760-Nicotinate-and-nicotinamide-metabolism sce01100-Metabolic-pathways sce01110-Biosynthesis-of-secondary-metabolites | 1 | 1 | -1 | 1 | 1 | 1 |
| DUTPDP | M-dutp-c M-h2o-c | M-dump-c M-h-c M-ppi-c | sce00240-Pyrimidine-metabolism sce01100-Metabolic-pathways | -1 | 1 | -1 | 1 | 0 | 1 |
| D-LACDm | M-ficytc-m M-lac–D-m | M-focytc-m M-pyr-m | sce00620-Pyruvate-metabolism sce01100-Metabolic-pathways | -1 | -1 | 1 | 1 | 1 | 1 |

| Reaction data | | | | Wild Type | | | $mip6\Delta$ | | |
|---|---|---|---|---|---|---|---|---|---|
| Rxn | Reactants | Products | Pathways | t0 | t20 | t120 | t0 | t20 | t120 |
| ECOAH11p | M-h2o-x M-hxc2coa-x | M-3hxccoa-x | sce00410-beta-Alanine-metabolism sce00640-Propanoate-metabolism sce01100-Metabolic-pathways sce01200-Carbon-metabolism | 1 | -1 | 1 | 1 | 1 | 1 |
| ECOAH4p | M-3hdcoa-x | M-dc2coa-x M-h2o-x | sce00410-beta-Alanine-metabolism sce00640-Propanoate-metabolism sce01100-Metabolic-pathways sce01200-Carbon-metabolism | 1 | -1 | 1 | 1 | 1 | 1 |
| ECOAH6p | M-3htdcoa-x | M-h2o-x M-td2coa-x | sce00410-beta-Alanine-metabolism sce00640-Propanoate-metabolism sce01100-Metabolic-pathways sce01200-Carbon-metabolism | 1 | -1 | 1 | 1 | 1 | 1 |
| ECOAH7p | M-3hhdcoa-x | M-h2o-x M-hdd2coa-x | sce00410-beta-Alanine-metabolism sce00640-Propanoate-metabolism sce01100-Metabolic-pathways sce01200-Carbon-metabolism | 1 | -1 | 1 | 1 | 1 | 1 |
| ECOAH8p | M-3hodcoa-x | M-h2o-x M-od2coa-x | sce00410-beta-Alanine-metabolism sce00640-Propanoate-metabolism sce01100-Metabolic-pathways sce01200-Carbon-metabolism | 1 | -1 | 1 | 1 | 1 | 1 |
| EX-2mbald-e | M-2mbald-e | NA | NA | 1 | 1 | -1 | 1 | 1 | 1 |
| EX-acald-e | M-acald-e | NA | NA | -1 | 1 | 1 | 1 | 1 | 1 |
| EX-asp–L-e | M-asp–L-e | NA | NA | 1 | 1 | -1 | -1 | -1 | 1 |
| EX-for-e | M-for-e | NA | NA | 1 | -1 | 1 | 1 | 1 | 1 |
| EX-fum-e | M-fum-e | NA | NA | -1 | 1 | 1 | 1 | 1 | 1 |
| EX-glu–L-e | M-glu–L-e | NA | NA | 1 | -1 | 1 | 1 | 1 | 1 |
| EX-glyc-e | M-glyc-e | NA | NA | 1 | -1 | 1 | -1 | 1 | -1 |
| EX-lac–D-e | M-lac–D-e | NA | NA | -1 | 1 | 1 | 1 | 0 | 1 |
| EX-nh4-e | M-nh4-e | NA | NA | -1 | 1 | -1 | 1 | 1 | 1 |
| EX-oaa-e | M-oaa-e | NA | NA | -1 | 1 | -1 | 1 | 1 | 1 |
| EX-orn-e | M-orn-e | NA | NA | -1 | -1 | -1 | 1 | 1 | 1 |
| EX-pyr-e | M-pyr-e | NA | NA | 1 | -1 | -1 | 1 | 1 | 1 |
| EX-succ-e | M-succ-e | NA | NA | -1 | 0 | 1 | 1 | 0 | -1 |
| FACOAL161 | M-atp-c M-coa-c M-hdcea-c | M-TM-atp-c M-amp-c M-hdcoa-c M-ppi-c | sce00061-Fatty-acid-biosynthesis sce00071-Fatty-acid-degradation sce01100-Metabolic-pathways sce01212-Fatty-acid-metabolism sce04146-Peroxisome | 1 | 1 | -1 | 1 | 1 | 1 |
| FACOAL80p | M-atp-x M-coa-x M-octa-x | M-amp-x M-occoa-x M-ppi-x | sce00061-Fatty-acid-biosynthesis sce00071-Fatty-acid-degradation sce01100-Metabolic-pathways sce01212-Fatty-acid-metabolism sce04146-Peroxisome | 1 | 1 | -1 | 1 | 0 | 1 |

| Reaction data | | | | Wild Type | | | $mip6\Delta$ | | |
|---|---|---|---|---|---|---|---|---|---|
| Rxn | Reactants | Products | Pathways | t0 | t20 | t120 | t0 | t20 | t120 |
| FALDH | M-fald-c M-gthrd-c M-nad-c | M-Sfglutth-c M-TM-gthrd-c M-TM-nad-c M-h-c M-nadh-c | sce00010-Glycolysis-/-Gluconeogenesis sce00071-Fatty-acid-degradation sce00350-Tyrosine-metabolism sce00680-Methane-metabolism sce01100-Metabolic-pathways sce01110-Biosynthesis-of-secondary-metabolites sce01200-Carbon-metabolism | -1 | 1 | 1 | 1 | 1 | 1 |
| FBP | M-fdp-c M-h2o-c | M-TM-fdp-c M-f6p-c M-pi-c | sce00010-Glycolysis-/-Gluconeogenesis sce00030-Pentose-phosphate-pathway sce00051-Fructose-and-mannose-metabolism sce00680-Methane-metabolism sce01100-Metabolic-pathways sce01110-Biosynthesis-of-secondary-metabolites sce01200-Carbon-metabolism | -1 | -1 | -1 | 1 | 1 | 1 |
| FECOSTt | M-fecost-e | M-fecost-c | sce02010-ABC-transporters | -1 | 1 | 1 | 1 | 0 | 1 |
| FRDm | M-fadh2-m M-fum-m | M-fad-m M-succ-m | NA | 1 | -1 | 1 | 1 | 1 | 1 |
| FTHFLi | M-atp-c M-for-c M-thf-c | M-10fthf-c M-TM-atp-c M-adp-c M-pi-c | sce00670-One-carbon-pool-by-folate sce01100-Metabolic-pathways | -1 | -1 | 1 | 1 | 1 | 1 |
| G3PCt | M-g3pc-e | M-TM-g3pc-c M-g3pc-c | NA | -1 | 1 | 1 | 1 | 1 | 1 |
| G3PD1ir | M-dhap-c M-h-c M-nadh-c | M-TM-nad-c M-glyc3p-c M-nad-c | sce00564-Glycerophospholipid-metabolism sce01110-Biosynthesis-of-secondary-metabolites sce04011-MAPK-signaling-pathway | 1 | -1 | 1 | 1 | 1 | 1 |
| G3PIt | M-g3pi-e | M-g3pi-c | NA | 1 | 1 | -1 | -1 | 1 | -1 |
| G3PT | M-glyc3p-c M-h2o-c | M-TM-glyc-c M-glyc-c M-pi-c | sce00561-Glycerolipid-metabolism sce01100-Metabolic-pathways | -1 | 1 | 1 | 1 | 1 | 1 |
| G5SADrm | M-glu5sa-m | M-1pyr5c-m M-h2o-m M-h-m | NA | 1 | -1 | 1 | 1 | 1 | 1 |
| G5SD2 | M-glu5p-c M-h-c M-nadh-c | M-TM-nad-c M-glu5sa-c M-nad-c M-pi-c | sce00330-Arginine-and-proline-metabolism sce00332-Carbapenem-biosynthesis sce01100-Metabolic-pathways sce01110-Biosynthesis-of-secondary-metabolites sce01230-Biosynthesis-of-amino-acids | 1 | -1 | 1 | 1 | 1 | 1 |

| Reaction data | | | | Wild Type | | | $mip6\Delta$ | | |
|---|---|---|---|---|---|---|---|---|---|
| Rxn | Reactants | Products | Pathways | t0 | t20 | t120 | t0 | t20 | t120 |
| G6PDH2r | M-g6p-c M-nadp-c | M-6pgl-c M-h-c M-nadph-c | sce00030-Pentose-phosphate-pathway sce00480-Glutathione-metabolism sce01100-Metabolic-pathways sce01110-Biosynthesis-of-secondary-metabolites sce01200-Carbon-metabolism | -1 | 0 | 1 | 1 | 1 | 1 |
| GCC2bim | M-alpam-m M-thf-m | M-dhlam-m M-mlthf-m M-nh4-m | sce00010-Glycolysis-/-Gluconeogenesis sce00020-Citrate-cycle-(TCA-cycle) sce00260-Glycine,-serine-and-threonine-metabolism sce00280-Valine,-leucine-and-isoleucine-degradation sce00310-Lysine-degradation sce00380-Tryptophan-metabolism sce00620-Pyruvate-metabolism sce00630-Glyoxylate-and-dicarboxylate-metabolism sce00640-Propanoate-metabolism sce00670-One-carbon-pool-by-folate sce01100-Metabolic-pathways sce01110-Biosynthesis-of-secondary-metabolites sce01200-Carbon-metabolism | 1 | -1 | 1 | 1 | 1 | 1 |
| GLCS2 | M-udpg-c | M-TM-udpg-c M-glycogen-c M-h-c M-udp-c | sce00500-Starch-and-sucrose-metabolism sce01100-Metabolic-pathways sce01110-Biosynthesis-of-secondary-metabolites | 1 | -1 | 1 | 1 | 1 | 1 |
| GLNt2r | M-gln–L-e M-h-e | M-TM-gln–L-c M-gln–L-c M-h-c | NA | -1 | -1 | 1 | 1 | 1 | 1 |
| GLUSx | M-akg-c M-gln–L-c M-h-c M-nadh-c | M-TM-gln–L-c M-TM-glu–L-c M-TM-nad-c M-glu–L-c M-nad-c | sce00250-Alanine,-aspartate-and-glutamate-metabolism sce00910-Nitrogen-metabolism sce01100-Metabolic-pathways sce01110-Biosynthesis-of-secondary-metabolites sce01230-Biosynthesis-of-amino-acids | -1 | -1 | 1 | 1 | 1 | 1 |
| GLUt7m | M-glu–L-c | M-TM-glu–L-c M-glu–L-m | NA | -1 | -1 | -1 | 1 | 1 | 1 |
| GLYCDy | M-glyc-c M-nadp-c | M-TM-glyc-c M-dha-c M-h-c M-nadph-c | sce00561-Glycerolipid-metabolism sce01100-Metabolic-pathways | 1 | -1 | 1 | 1 | 1 | 1 |

| Reaction data | | | | Wild Type | | | $mip6\Delta$ | | |
|---|---|---|---|---|---|---|---|---|---|
| Rxn | Reactants | Products | Pathways | t0 | t20 | t120 | t0 | t20 | t120 |
| GLYK | M-atp-c M-glyc-c | M-TM-atp-c M-TM-glyc-c M-adp-c M-glyc3p-c M-h-c | sce00561-Glycerolipid-metabolism sce01100-Metabolic-pathways | 1 | 1 | -1 | 1 | 1 | 1 |
| GLYOX | M-h2o-c M-lgt–S-c | M-TM-gthrd-c M-gthrd-c M-h-c M-lac–D-c | sce00620-Pyruvate-metabolism sce01100-Metabolic-pathways | 1 | -1 | -1 | 1 | 0 | 1 |
| GNNUC | M-gsn-c M-h2o-c | M-gua-c M-rib–D-c | sce00240-Pyrimidine-metabolism sce00760-Nicotinate-and-nicotinamide-metabolism sce01100-Metabolic-pathways | 1 | 1 | -1 | 1 | 1 | 1 |
| H2Otp | M-h2o-c | M-h2o-x | NA | -1 | 0 | 1 | 1 | 1 | 1 |
| HACD10p | M-3hxccoa-x M-nad-x | M-3ohxccoa-x M-h-x M-nadh-x | sce00410-beta-Alanine-metabolism sce00640-Propanoate-metabolism sce01100-Metabolic-pathways sce01200-Carbon-metabolism | -1 | -1 | -1 | 1 | 1 | -1 |
| HACD7p | M-3ohdcoa-x M-h-x M-nadh-x | M-3hhdcoa-x M-nad-x | sce00410-beta-Alanine-metabolism sce00640-Propanoate-metabolism sce01100-Metabolic-pathways sce01200-Carbon-metabolism | -1 | -1 | -1 | 1 | 1 | -1 |
| HCO3E | M-co2-c M-h2o-c | M-h-c M-hco3-c | NA | 1 | 1 | -1 | -1 | 0 | -1 |
| HCO3tn | M-hco3-c | M-hco3-n | NA | -1 | 1 | -1 | 1 | 1 | 1 |
| HMGCOAS | M-coa-c M-h-c M-hmgcoa-c | M-aacoa-c M-accoa-c M-h2o-c | sce00072-Synthesis-and-degradation-of-ketone-bodies sce00280-Valine,-leucine-and-isoleucine-degradation sce00650-Butanoate-metabolism sce00900-Terpenoid-backbone-biosynthesis sce01100-Metabolic-pathways sce01110-Biosynthesis-of-secondary-metabolites | -1 | -1 | 1 | 1 | 1 | 1 |
| HSDxi | M-aspsa-c M-h-c M-nadh-c | M-TM-nad-c M-hom–L-c M-nad-c | sce00260-Glycine,-serine-and-threonine-metabolism sce00270-Cysteine-and-methionine-metabolism sce00300-Lysine-biosynthesis sce01100-Metabolic-pathways sce01110-Biosynthesis-of-secondary-metabolites sce01230-Biosynthesis-of-amino-acids | 1 | 1 | -1 | 1 | 1 | 1 |

| Reaction data | | | | Wild Type | | | $mip6\Delta$ | | |
|---|---|---|---|---|---|---|---|---|---|
| Rxn | Reactants | Products | Pathways | t0 | t20 | t120 | t0 | t20 | t120 |
| ICL | M-icit-c | M-TM-succ-c M-glx-c M-succ-c | sce00630-Glyoxylate-and-dicarboxylate-metabolism sce01100-Metabolic-pathways sce01110-Biosynthesis-of-secondary-metabolites sce01200-Carbon-metabolism | -1 | 1 | 1 | 1 | 1 | 1 |
| ILEt2r | M-h-e M-ile–L-e | M-TM-ile–L-c M-h-c M-ile–L-c | NA | -1 | 1 | -1 | 1 | 1 | 1 |
| ILEtmi | M-ile–L-m | M-TM-ile–L-c M-ile–L-c | NA | -1 | 1 | -1 | 1 | 1 | 1 |
| IPPMIb | M-2ippm-c M-h2o-c | M-3c3hmp-c | sce00290-Valine,-leucine-and-isoleucine-biosynthesis sce01100-Metabolic-pathways sce01110-Biosynthesis-of-secondary-metabolites sce01210-2-Oxocarboxylic-acid-metabolism sce01230-Biosynthesis-of-amino-acids | 1 | -1 | 1 | 1 | 1 | 1 |
| IPPSm | M-3mob-m M-accoa-m M-h2o-m | M-3c3hmp-m M-coa-m M-h-m | sce00290-Valine,-leucine-and-isoleucine-biosynthesis sce00620-Pyruvate-metabolism sce01100-Metabolic-pathways sce01110-Biosynthesis-of-secondary-metabolites sce01210-2-Oxocarboxylic-acid-metabolism sce01230-Biosynthesis-of-amino-acids | -1 | -1 | 1 | 1 | 1 | 1 |
| ITCOALm | M-atp-m M-coa-m M-itacon-m | M-adp-m M-itaccoa-m M-pi-m | sce00020-Citrate-cycle-(TCA-cycle) sce00640-Propanoate-metabolism sce01100-Metabolic-pathways sce01110-Biosynthesis-of-secondary-metabolites sce01200-Carbon-metabolism | -1 | -1 | 1 | 1 | 1 | 1 |
| LALDO3 | M-h-c M-mthgxl-c M-nadph-c | M-lald–L-c M-nadp-c | sce00040-Pentose-and-glucuronate-interconversions sce00620-Pyruvate-metabolism sce00640-Propanoate-metabolism sce01100-Metabolic-pathways sce04011-MAPK-signaling-pathway | -1 | 1 | 1 | 1 | 1 | 1 |
| LEUt2r | M-h-e M-leu–L-e | M-TM-leu–L-c M-h-c M-leu–L-c | NA | 1 | 0 | -1 | 1 | 1 | 1 |
| LGTHL | M-gthrd-c M-mthgxl-c | M-TM-gthrd-c M-lgt–S-c | sce00620-Pyruvate-metabolism sce01100-Metabolic-pathways | 1 | -1 | -1 | 1 | 0 | 1 |

| Reaction data | | | | Wild Type | | | $mip6\Delta$ | | |
|---|---|---|---|---|---|---|---|---|---|
| Rxn | Reactants | Products | Pathways | t0 | t20 | t120 | t0 | t20 | t120 |
| LNS14DM | M-h-c M-lanost-c M-nadph-c M-o2-c | M-44mctr-c M-for-c M-h2o-c M-nadp-c | sce00100-Steroid-biosynthesis sce01100-Metabolic-pathways sce01110-Biosynthesis-of-secondary-metabolites | 1 | -1 | 1 | 1 | 1 | 1 |
| LPP-SC | M-dagpy-SC-c M-h2o-c | M-h-c M-pa-SC-c M-pi-c | sce00561-Glycerolipid-metabolism sce00564-Glycerophospholipid-metabolism sce01110-Biosynthesis-of-secondary-metabolites | 1 | 1 | -1 | 1 | 1 | 1 |
| LSERDHr | M-nadp-c M-ser–L-c | M-2amsa-c M-TM-ser–L-c M-h-c M-nadph-c | sce00240-Pyrimidine-metabolism sce00260-Glycine,-serine-and-threonine-metabolism sce01100-Metabolic-pathways | 1 | -1 | -1 | 1 | 1 | -1 |
| MALt2r | M-h-e M-mal–L-e | M-h-c M-mal–L-c | NA | -1 | 1 | -1 | -1 | -1 | -1 |
| MDHm | M-mal–L-m M-nad-m | M-h-m M-nadh-m M-oaa-m | sce00020-Citrate-cycle-(TCA-cycle) sce00270-Cysteine-and-methionine-metabolism sce00620-Pyruvate-metabolism sce00630-Glyoxylate-and-dicarboxylate-metabolism sce01100-Metabolic-pathways sce01110-Biosynthesis-of-secondary-metabolites sce01200-Carbon-metabolism | -1 | -1 | 1 | 1 | 1 | 1 |
| MEVK1 | M-atp-c M-mev–R-c | M-5pmev-c M-TM-atp-c M-adp-c M-h-c | sce00900-Terpenoid-backbone-biosynthesis sce01100-Metabolic-pathways sce01110-Biosynthesis-of-secondary-metabolites sce04146-Peroxisome | -1 | 1 | -1 | 1 | 1 | 1 |
| MEVK3 | M-gtp-c M-mev–R-c | M-5pmev-c M-gdp-c M-h-c | sce00900-Terpenoid-backbone-biosynthesis sce01100-Metabolic-pathways sce01110-Biosynthesis-of-secondary-metabolites sce04146-Peroxisome | 1 | -1 | -1 | 1 | 1 | 1 |
| MEVK4 | M-mev–R-c M-utp-c | M-5pmev-c M-h-c M-udp-c | sce00900-Terpenoid-backbone-biosynthesis sce01100-Metabolic-pathways sce01110-Biosynthesis-of-secondary-metabolites sce04146-Peroxisome | 1 | -1 | 1 | 1 | 1 | 1 |
| NADPPPS | M-h2o-c M-nadp-c | M-TM-nad-c M-nad-c M-pi-c | NA | 1 | -1 | -1 | -1 | 1 | -1 |

| Reaction data | | | | Wild Type | | | $mip6\Delta$ | | |
|---|---|---|---|---|---|---|---|---|---|
| Rxn | Reactants | Products | Pathways | t0 | t20 | t120 | t0 | t20 | t120 |
| NDP3 | M-gdp-c M-h2o-c | M-TM-gmp-c M-gmp-c M-h-c M-pi-c | sce00230-Purine-metabolism sce00240-Pyrimidine-metabolism sce01100-Metabolic-pathways | 1 | 1 | -1 | 1 | 1 | 1 |
| NDP7 | M-h2o-c M-udp-c | M-h-c M-pi-c M-ump-c | sce00230-Purine-metabolism sce00240-Pyrimidine-metabolism sce01100-Metabolic-pathways | -1 | -1 | 1 | 1 | 1 | 1 |
| NDPK3 | M-atp-c M-cdp-c | M-TM-atp-c M-adp-c M-ctp-c | sce00230-Purine-metabolism sce00240-Pyrimidine-metabolism sce01100-Metabolic-pathways sce01110-Biosynthesis-of-secondary-metabolites | -1 | -1 | 1 | 1 | 1 | 1 |
| NDPK8 | M-atp-c M-dadp-c | M-TM-atp-c M-adp-c M-datp-c | sce00230-Purine-metabolism sce00240-Pyrimidine-metabolism sce01100-Metabolic-pathways sce01110-Biosynthesis-of-secondary-metabolites | -1 | -1 | 1 | 1 | 1 | 1 |
| NH4tm | M-nh4-c | M-nh4-m | NA | -1 | 1 | 1 | 1 | 1 | 1 |
| NTD2 | M-h2o-c M-ump-c | M-pi-c M-uri-c | sce00760-Nicotinate-and-nicotinamide-metabolism | -1 | -1 | -1 | 1 | 1 | 1 |
| NTP3 | M-gtp-c M-h2o-c | M-gdp-c M-h-c M-pi-c | NA | -1 | -1 | 1 | -1 | -1 | -1 |
| OAAt | M-oaa-c | M-oaa-e | NA | -1 | 1 | -1 | 1 | 1 | 1 |
| OHPBAT | M-glu–L-c M-ohpb-c | M-TM-glu–L-c M-akg-c M-phthr-c | sce00260-Glycine,-serine-and-threonine-metabolism sce00270-Cysteine-and-methionine-metabolism sce00680-Methane-metabolism sce00750-Vitamin-B6-metabolism sce01100-Metabolic-pathways sce01110-Biosynthesis-of-secondary-metabolites sce01200-Carbon-metabolism sce01230-Biosynthesis-of-amino-acids | -1 | 0 | -1 | 1 | 1 | 1 |
| ORNTA | M-akg-c M-orn-c | M-TM-glu–L-c M-glu5sa-c M-glu–L-c | sce00330-Arginine-and-proline-metabolism sce01100-Metabolic-pathways sce01110-Biosynthesis-of-secondary-metabolites | -1 | 0 | -1 | 1 | 1 | 1 |
| P5CDm | M-1pyr5c-m M-h2o-m M-nad-m | M-glu–L-m M-h-m M-nadh-m | NA | 1 | -1 | 1 | -1 | 1 | 1 |
| PAK-SC | M-atp-c M-pa-SC-c | M-TM-atp-c M-adp-c M-dagpy-SC-c | NA | 1 | 1 | -1 | 1 | 1 | 1 |
| PAPtm | M-pap-c | M-pap-m | NA | 1 | 1 | -1 | -1 | -1 | -1 |
| PDE1 | M-camp-c M-h2o-c | M-amp-c M-h-c | sce00230-Purine-metabolism sce01100-Metabolic-pathways | 1 | -1 | -1 | 1 | 1 | 1 |

| Reaction data | | | | Wild Type | | | $mip6\Delta$ | | |
|---|---|---|---|---|---|---|---|---|---|
| Rxn | Reactants | Products | Pathways | t0 | t20 | t120 | t0 | t20 | t120 |
| GCC2cm-copy2 | M-dhlam-m M-nad-m | M-h-m M-lpam-m M-nadh-m | sce00010-Glycolysis-/-Gluconeogenesis sce00020-Citrate-cycle-(TCA-cycle) sce00260-Glycine,-serine-and-threonine-metabolism sce00280-Valine,-leucine-and-isoleucine-degradation sce00310-Lysine-degradation sce00380-Tryptophan-metabolism sce00620-Pyruvate-metabolism sce00630-Glyoxylate-and-dicarboxylate-metabolism sce00640-Propanoate-metabolism sce01100-Metabolic-pathways sce01110-Biosynthesis-of-secondary-metabolites sce01200-Carbon-metabolism | -1 | -1 | 1 | 1 | 1 | 1 |
| PEtm-SC | M-pe-SC-c | M-pe-SC-m | NA | -1 | 1 | 1 | 1 | 1 | 1 |
| PGI | M-g6p-c | M-f6p-c | sce00010-Glycolysis-/-Gluconeogenesis sce00030-Pentose-phosphate-pathway sce00500-Starch-and-sucrose-metabolism sce00520-Amino-sugar-and-nucleotide-sugar-metabolism sce01100-Metabolic-pathways sce01110-Biosynthesis-of-secondary-metabolites sce01200-Carbon-metabolism | -1 | -1 | 1 | 1 | 1 | 1 |
| PGM | M-2pg-c | M-3pg-c | sce00010-Glycolysis-/-Gluconeogenesis sce00260-Glycine,-serine-and-threonine-metabolism sce00680-Methane-metabolism sce01100-Metabolic-pathways sce01110-Biosynthesis-of-secondary-metabolites sce01200-Carbon-metabolism sce01230-Biosynthesis-of-amino-acids | 1 | 1 | -1 | 1 | 1 | 1 |
| PI35BP5P-SC | M-h2o-c M-ptd135bp-SC-c | M-pi-c M-ptd3ino-SC-c | sce00562-Inositol-phosphate-metabolism sce01100-Metabolic-pathways sce04070-Phosphatidylinositol-signaling-system | -1 | 1 | 1 | 1 | 1 | 1 |

| Reaction data | | | | Wild Type | | | $mip6\Delta$ | | |
|---|---|---|---|---|---|---|---|---|---|
| Rxn | Reactants | Products | Pathways | t0 | t20 | t120 | t0 | t20 | t120 |
| PI3P5K-SC | M-atp-c M-ptd3ino-SC-c | M-TM-atp-c M-adp-c M-h-c M-ptd135bp-SC-c | sce00562-Inositol-phosphate-metabolism sce01100-Metabolic-pathways sce04070-Phosphatidylinositol-signaling-system sce04145-Phagosome | -1 | 1 | 1 | 1 | 1 | 1 |
| PLD-SC | M-h2o-c M-pc-SC-c | M-chol-c M-h-c M-pa-SC-c | sce00564-Glycerophospholipid-metabolism sce00565-Ether-lipid-metabolism sce01100-Metabolic-pathways sce01110-Biosynthesis-of-secondary-metabolites sce04144-Endocytosis | -1 | -1 | 1 | 1 | 1 | 1 |
| PPND | M-nad-c M-pphn-c | M-34hpp-c M-TM-nad-c M-co2-c M-nadh-c | NA | 1 | 1 | 1 | 1 | -1 | 1 |
| PROt2r | M-h-e M-pro–L-e | M-TM-pro–L-c M-h-c M-pro–L-c | NA | -1 | -1 | 1 | 1 | 1 | 1 |
| PSERSm-SC | M-cdpdag-SC-m M-ser–L-m | M-cmp-m M-h-m M-ps-SC-m | sce00260-Glycine,-serine-and-threonine-metabolism sce00564-Glycerophospholipid-metabolism sce01100-Metabolic-pathways sce01110-Biosynthesis-of-secondary-metabolites | -1 | 1 | -1 | 1 | 0 | 1 |
| PTRCt3i | M-h-c M-ptrc-e | M-h-e M-ptrc-c | NA | -1 | -1 | 1 | 1 | 1 | 1 |
| PTRCtex2 | M-ptrc-c | M-ptrc-e | NA | -1 | 1 | 1 | 1 | 1 | 1 |
| PUNP1 | M-adn-c M-pi-c | M-ade-c M-r1p-c | sce00230-Purine-metabolism sce00240-Pyrimidine-metabolism sce00760-Nicotinate-and-nicotinamide-metabolism sce01100-Metabolic-pathways sce01110-Biosynthesis-of-secondary-metabolites | 1 | -1 | -1 | 1 | 1 | 1 |
| PUNP6 | M-din-c M-pi-c | M-2dr1p-c M-hxan-c | sce00230-Purine-metabolism sce00240-Pyrimidine-metabolism sce00760-Nicotinate-and-nicotinamide-metabolism sce01100-Metabolic-pathways sce01110-Biosynthesis-of-secondary-metabolites | -1 | -1 | 1 | 1 | 1 | 1 |
| PYRt2 | M-h-e M-pyr-e | M-h-c M-pyr-c | NA | -1 | 0 | 1 | 1 | 1 | 1 |
| RNDR3 | M-cdp-c M-trdrd-c | M-dcdp-c M-h2o-c M-trdox-c | sce00230-Purine-metabolism sce00240-Pyrimidine-metabolism sce00480-Glutathione-metabolism sce01100-Metabolic-pathways | 1 | 1 | -1 | 1 | 1 | 1 |
| SERAT | M-accoa-c M-ser–L-c | M-TM-ser–L-c M-acser-c M-coa-c | NA | 1 | -1 | 1 | 1 | 1 | 1 |

| Reaction data | | | | Wild Type | | | $mip6\Delta$ | | |
|---|---|---|---|---|---|---|---|---|---|
| Rxn | Reactants | Products | Pathways | t0 | t20 | t120 | t0 | t20 | t120 |
| SLFAT | M-adp-c M-h-c M-so4-c | M-aps-c M-pi-c | sce00230-Purine-metabolism sce00920-Sulfur-metabolism sce01100-Metabolic-pathways | 1 | 1 | -1 | 1 | 1 | 1 |
| THIORDXi | M-h2o2-c M-trdrd-c | M-h2o-c M-trdox-c | NA | -1 | -1 | 1 | 1 | 1 | 1 |
| THIORDXp | M-h2o2-x M-trdrd-x | M-h2o-x M-trdox-x | sce04122-Sulfur-relay-system | -1 | -1 | -1 | 1 | 1 | 1 |
| THMP | M-h2o-c M-thmmp-c | M-TM-thm-c M-pi-c M-thm-c | NA | 1 | -1 | -1 | 1 | 1 | 1 |
| THRt2r | M-h-e M-thr–L-e | M-h-c M-thr–L-c | NA | -1 | -1 | 1 | 1 | 1 | 1 |
| TKT1 | M-r5p-c M-xu5p–D-c | M-g3p-c M-s7p-c | sce00030-Pentose-phosphate-pathway sce01100-Metabolic-pathways sce01110-Biosynthesis-of-secondary-metabolites sce01200-Carbon-metabolism sce01230-Biosynthesis-of-amino-acids | 1 | 1 | -1 | 1 | 1 | 1 |
| TMDK1 | M-atp-c M-thymd-c | M-TM-atp-c M-adp-c M-dtmp-c M-h-c | NA | -1 | 1 | 1 | 1 | -1 | 1 |
| TMN | M-h2o-c M-thm-c | M-4ahmmp-c M-4mhetz-c M-TM-thm-c M-h-c | NA | 1 | 1 | -1 | 1 | 1 | 1 |
| TREH | M-h2o-c M-tre-c | M-TM-glc–D-c M-TM-tre-c M-glc–D-c | sce00500-Starch-and-sucrose-metabolism sce01100-Metabolic-pathways sce01110-Biosynthesis-of-secondary-metabolites | 1 | -1 | -1 | 1 | 1 | 1 |
| TREt2v | M-h-c M-tre-c | M-TM-tre-c M-h-v M-tre-v | NA | 1 | 1 | -1 | 1 | 1 | 1 |
| UPPRT | M-prpp-c M-ura-c | M-ppi-c M-ump-c | sce00240-Pyrimidine-metabolism sce01100-Metabolic-pathways | 1 | 1 | -1 | 1 | 0 | 1 |
| UREASE | M-atp-c M-hco3-c M-urea-c | M-TM-atp-c M-adp-c M-allphn-c M-h-c M-pi-c | sce00220-Arginine-biosynthesis sce00791-Atrazine-degradation sce01100-Metabolic-pathways | -1 | -1 | -1 | 1 | 1 | 1 |
| URIK1 | M-atp-c M-uri-c | M-TM-atp-c M-adp-c M-h-c M-ump-c | sce00240-Pyrimidine-metabolism sce01100-Metabolic-pathways | -1 | 1 | -1 | 1 | 1 | 1 |
| VALt2r | M-h-e M-val–L-e | M-TM-val–L-c M-h-c M-val–L-c | NA | 1 | -1 | -1 | 1 | 0 | 1 |
| 2MBALDt-reverse | M-2mbald-e | M-2mbald-c | NA | 1 | -1 | 1 | 1 | 1 | 1 |
| 3C3HMPtm-reverse | M-3c3hmp-m | M-3c3hmp-c | NA | -1 | -1 | 1 | 1 | 1 | 1 |
| ACALDt-reverse | M-acald-c | M-acald-e | NA | -1 | 1 | 1 | 1 | 1 | 1 |
| ACALDtm-reverse | M-acald-c | M-acald-m | NA | -1 | 1 | -1 | 1 | 1 | 1 |

| Reaction data | | | | Wild Type | | | $mip6\Delta$ | | |
|---|---|---|---|---|---|---|---|---|---|
| Rxn | Reactants | Products | Pathways | t0 | t20 | t120 | t0 | t20 | t120 |
| ACCOAC-reverse | M-adp-c M-h-c M-malcoa-c M-pi-c | M-TM-atp-c M-accoa-c M-atp-c M-hco3-c | sce00061-Fatty-acid-biosynthesis sce00620-Pyruvate-metabolism sce00640-Propanoate-metabolism sce01100-Metabolic-pathways sce01110-Biosynthesis-of-secondary-metabolites sce01212-Fatty-acid-metabolism | -1 | 1 | 1 | 1 | 1 | 1 |
| ADK1-reverse | M-adp-c | M-TM-atp-c M-amp-c M-atp-c | sce00230-Purine-metabolism sce00730-Thiamine-metabolism sce01100-Metabolic-pathways sce01110-Biosynthesis-of-secondary-metabolites | -1 | -1 | -1 | 1 | 1 | 1 |
| ADK3m-reverse | M-adp-m M-gdp-m | M-amp-m M-gtp-m | sce00230-Purine-metabolism sce00730-Thiamine-metabolism sce01100-Metabolic-pathways sce01110-Biosynthesis-of-secondary-metabolites | -1 | 1 | -1 | 1 | 1 | 1 |
| ADK4m-reverse | M-adp-m M-idp-m | M-amp-m M-itp-m | sce00230-Purine-metabolism sce00730-Thiamine-metabolism sce01100-Metabolic-pathways sce01110-Biosynthesis-of-secondary-metabolites | -1 | -1 | 1 | 1 | 1 | 1 |
| AKGMAL-reverse | M-akg-e M-mal–L-c | M-akg-c M-mal–L-e | NA | -1 | -1 | 1 | 1 | 1 | 1 |
| ASPt2n-reverse | M-asp–L-n M-h-n | M-TM-asp–L-c M-asp–L-c M-h-c | NA | 1 | -1 | 1 | 1 | 1 | 1 |
| ASPt2r-reverse | M-asp–L-c M-h-c | M-TM-asp–L-c M-asp–L-e M-h-e | NA | 1 | 1 | -1 | 1 | 1 | 1 |
| BTDD-RR-reverse | M-actn–R-c M-h-c M-nadh-c | M-TM-nad-c M-btd-RR-c M-nad-c | sce00650-Butanoate-metabolism | -1 | -1 | -1 | -1 | 0 | 1 |
| CYSt2r-reverse | M-cys–L-c M-h-c | M-cys–L-e M-h-e | NA | -1 | 1 | -1 | 1 | 1 | 1 |
| CYTK1-reverse | M-adp-c M-cdp-c | M-TM-atp-c M-TM-cmp-c M-atp-c M-cmp-c | NA | -1 | -1 | -1 | -1 | 0 | 1 |
| DASYN-SC-reverse | M-cdpdag-SC-c M-ppi-c | M-ctp-c M-h-c M-pa-SC-c | sce00564-Glycerophospholipid-metabolism sce01100-Metabolic-pathways sce01110-Biosynthesis-of-secondary-metabolites sce04070-Phosphatidylinositol-signaling-system | 1 | 1 | -1 | 1 | 1 | 1 |

| Reaction data | | | | Wild Type | | | $mip6\Delta$ | | |
|---|---|---|---|---|---|---|---|---|---|
| Rxn | Reactants | Products | Pathways | t0 | t20 | t120 | t0 | t20 | t120 |
| DURIPP-reverse | M-2dr1p-c M-ura-c | M-duri-c M-pi-c | sce00230-Purine-metabolism sce00240-Pyrimidine-metabolism sce00760-Nicotinate-and-nicotinamide-metabolism sce01100-Metabolic-pathways sce01110-Biosynthesis-of-secondary-metabolites | -1 | 1 | 1 | 1 | 1 | 1 |
| ECOAH11p-reverse | M-3hxccoa-x | M-h2o-x M-hxc2coa-x | sce00410-beta-Alanine-metabolism sce00640-Propanoate-metabolism sce01100-Metabolic-pathways sce01200-Carbon-metabolism | 1 | -1 | 1 | 1 | 1 | 1 |
| ECOAH4p-reverse | M-dc2coa-x M-h2o-x | M-3hdcoa-x | sce00410-beta-Alanine-metabolism sce00640-Propanoate-metabolism sce01100-Metabolic-pathways sce01200-Carbon-metabolism | 1 | -1 | 1 | 1 | 1 | 1 |
| ECOAH6p-reverse | M-h2o-x M-td2coa-x | M-3htdcoa-x | sce00410-beta-Alanine-metabolism sce00640-Propanoate-metabolism sce01100-Metabolic-pathways sce01200-Carbon-metabolism | 1 | -1 | 1 | 1 | 1 | 1 |
| ECOAH7p-reverse | M-h2o-x M-hdd2coa-x | M-3hhdcoa-x | sce00410-beta-Alanine-metabolism sce00640-Propanoate-metabolism sce01100-Metabolic-pathways sce01200-Carbon-metabolism | 1 | -1 | 1 | 1 | 1 | 1 |
| ECOAH8p-reverse | M-h2o-x M-od2coa-x | M-3hodcoa-x | sce00410-beta-Alanine-metabolism sce00640-Propanoate-metabolism sce01100-Metabolic-pathways sce01200-Carbon-metabolism | 1 | -1 | 1 | 1 | 1 | 1 |
| ENO-reverse | M-h2o-c M-pep-c | M-2pg-c | sce00010-Glycolysis-/-Gluconeogenesis sce00680-Methane-metabolism sce01100-Metabolic-pathways sce01110-Biosynthesis-of-secondary-metabolites sce01200-Carbon-metabolism sce01230-Biosynthesis-of-amino-acids sce03018-RNA-degradation | 1 | 1 | -1 | 1 | 1 | 1 |
| EX-h-e-reverse | NA | M-h-e | NA | -1 | -1 | 1 | 1 | 1 | 1 |

| Reaction data | | | | Wild Type | | | $mip6\Delta$ | | |
|---|---|---|---|---|---|---|---|---|---|
| Rxn | Reactants | Products | Pathways | t0 | t20 | t120 | t0 | t20 | t120 |
| FACOAL140-reverse | M-amp-c M-ppi-c M-tdcoa-c | M-TM-atp-c M-atp-c M-coa-c M-ttdca-c | sce00061-Fatty-acid-biosynthesis sce00071-Fatty-acid-degradation sce01100-Metabolic-pathways sce01212-Fatty-acid-metabolism sce04146-Peroxisome | -1 | 1 | 1 | 1 | 1 | 1 |
| FACOAL181-reverse | M-amp-c M-odecoa-c M-ppi-c | M-TM-atp-c M-atp-c M-coa-c M-ocdcea-c | sce00061-Fatty-acid-biosynthesis sce00071-Fatty-acid-degradation sce01100-Metabolic-pathways sce01212-Fatty-acid-metabolism sce04146-Peroxisome | 1 | -1 | 1 | 1 | 1 | 1 |
| FACOAL80p-reverse | M-amp-x M-occoa-x M-ppi-x | M-atp-x M-coa-x M-octa-x | sce00061-Fatty-acid-biosynthesis sce00071-Fatty-acid-degradation sce01100-Metabolic-pathways sce01212-Fatty-acid-metabolism sce04146-Peroxisome | 1 | 1 | -1 | 1 | 0 | 1 |
| FALDH-reverse | M-Sfglutth-c M-h-c M-nadh-c | M-TM-gthrd-c M-TM-nad-c M-fald-c M-gthrd-c M-nad-c | sce00010-Glycolysis-/-Gluconeogenesis sce00071-Fatty-acid-degradation sce00350-Tyrosine-metabolism sce00680-Methane-metabolism sce01100-Metabolic-pathways sce01110-Biosynthesis-of-secondary-metabolites sce01200-Carbon-metabolism | -1 | 1 | 1 | 1 | 1 | 1 |
| FBA-reverse | M-dhap-c M-g3p-c | M-TM-fdp-c M-fdp-c | sce00010-Glycolysis-/-Gluconeogenesis sce00030-Pentose-phosphate-pathway sce00051-Fructose-and-mannose-metabolism sce00680-Methane-metabolism sce01100-Metabolic-pathways sce01110-Biosynthesis-of-secondary-metabolites sce01200-Carbon-metabolism sce01230-Biosynthesis-of-amino-acids | 1 | -1 | 1 | 1 | 1 | 1 |
| FECOSTt-reverse | M-fecost-c | M-fecost-e | sce02010-ABC-transporters | -1 | 1 | 1 | 1 | 0 | 1 |
| FUMm-reverse | M-mal–L-m | M-fum-m M-h2o-m | sce00020-Citrate-cycle-(TCA-cycle) sce00620-Pyruvate-metabolism sce01100-Metabolic-pathways sce01110-Biosynthesis-of-secondary-metabolites sce01200-Carbon-metabolism | -1 | -1 | -1 | 1 | 1 | 1 |
| FUMt2r-reverse | M-fum-c M-h-c | M-fum-e M-h-e | NA | -1 | 1 | 1 | 1 | 1 | 1 |
| G3PCt-reverse | M-g3pc-c | M-TM-g3pc-c M-g3pc-e | NA | -1 | 1 | 1 | 1 | 1 | 1 |
| G3PIt-reverse | M-g3pi-c | M-g3pi-e | NA | 1 | 1 | -1 | -1 | 1 | -1 |

| Reaction data | | | | Wild Type | | | $mip6\Delta$ | | |
|---|---|---|---|---|---|---|---|---|---|
| Rxn | Reactants | Products | Pathways | t0 | t20 | t120 | t0 | t20 | t120 |
| GLUt2r-reverse | M-glu–L-c M-h-c | M-TM-glu–L-c M-glu–L-e M-h-e | NA | 1 | 1 | -1 | 1 | 1 | 1 |
| GLYCt-reverse | M-glyc-e | M-TM-glyc-c M-glyc-c | NA | -1 | 1 | -1 | 1 | 1 | 1 |
| H2Otp-reverse | M-h2o-x | M-h2o-c | NA | -1 | -1 | 1 | 1 | 1 | 1 |
| HACD10p-reverse | M-3ohxccoa-x M-h-x M-nadh-x | M-3hxccoa-x M-nad-x | sce00410-beta-Alanine-metabolism sce00640-Propanoate-metabolism sce01100-Metabolic-pathways sce01200-Carbon-metabolism | -1 | -1 | -1 | 1 | 1 | -1 |
| HACD7p-reverse | M-3hhdcoa-x M-nad-x | M-3ohdcoa-x M-h-x M-nadh-x | sce00410-beta-Alanine-metabolism sce00640-Propanoate-metabolism sce01100-Metabolic-pathways sce01200-Carbon-metabolism | -1 | -1 | -1 | 1 | 1 | -1 |
| HMGCOAtm-reverse | M-hmgcoa-m | M-hmgcoa-c | NA | 1 | 1 | -1 | 1 | 0 | 1 |
| ILETAm-reverse | M-3mop-m M-glu–L-m | M-akg-m M-ile–L-m | sce00270-Cysteine-and-methionine-metabolism sce00280-Valine,-leucine-and-isoleucine-degradation sce00290-Valine,-leucine-and-isoleucine-biosynthesis sce00770-Pantothenate-and-CoA-biosynthesis sce01100-Metabolic-pathways sce01110-Biosynthesis-of-secondary-metabolites sce01210-2-Oxocarboxylic-acid-metabolism sce01230-Biosynthesis-of-amino-acids | -1 | 1 | -1 | 1 | 1 | 1 |
| ITCOALm-reverse | M-adp-m M-itaccoa-m M-pi-m | M-atp-m M-coa-m M-itacon-m | sce00020-Citrate-cycle-(TCA-cycle) sce00640-Propanoate-metabolism sce01100-Metabolic-pathways sce01110-Biosynthesis-of-secondary-metabolites sce01200-Carbon-metabolism | -1 | -1 | 1 | 1 | 1 | 1 |
| LSERDHr-reverse | M-2amsa-c M-h-c M-nadph-c | M-TM-ser–L-c M-nadp-c M-ser–L-c | sce00240-Pyrimidine-metabolism sce00260-Glycine,-serine-and-threonine-metabolism sce01100-Metabolic-pathways | 1 | -1 | -1 | 1 | 1 | -1 |
| LYSt2r-reverse | M-h-c M-lys–L-c | M-TM-lys–L-c M-h-e M-lys–L-e | NA | -1 | -1 | 1 | 1 | 1 | 1 |
| L-LACtm-reverse | M-h-m M-lac–L-m | M-TM-lac–L-c M-h-c M-lac–L-c | NA | -1 | 1 | -1 | -1 | -1 | 1 |

| Reaction data | | | | Wild Type | | | $mip6\Delta$ | | |
|---|---|---|---|---|---|---|---|---|---|
| Rxn | Reactants | Products | Pathways | t0 | t20 | t120 | t0 | t20 | t120 |
| OHPBAT-reverse | M-akg-c M-phthr-c | M-TM-glu–L-c M-glu–L-c M-ohpb-c | sce00260-Glycine,-serine-and-threonine-metabolism sce00270-Cysteine-and-methionine-metabolism sce00680-Methane-metabolism sce00750-Vitamin-B6-metabolism sce01100-Metabolic-pathways sce01110-Biosynthesis-of-secondary-metabolites sce01200-Carbon-metabolism sce01230-Biosynthesis-of-amino-acids | -1 | 0 | -1 | 1 | 1 | 1 |
| PHEt2r-reverse | M-h-c M-phe–L-c | M-TM-phe–L-c M-h-e M-phe–L-e | NA | -1 | -1 | 1 | 1 | 1 | 1 |
| PPM-reverse | M-r5p-c | M-r1p-c | sce00010-Glycolysis-/-Gluconeogenesis sce00030-Pentose-phosphate-pathway sce00052-Galactose-metabolism sce00230-Purine-metabolism sce00500-Starch-and-sucrose-metabolism sce00520-Amino-sugar-and-nucleotide-sugar-metabolism sce01100-Metabolic-pathways sce01110-Biosynthesis-of-secondary-metabolites | -1 | -1 | -1 | 1 | 1 | 1 |
| PRASCSi-reverse | M-25aics-c M-adp-c M-h-c M-pi-c | M-5aizc-c M-TM-asp–L-c M-TM-atp-c M-asp–L-c M-atp-c | sce00230-Purine-metabolism sce01100-Metabolic-pathways sce01110-Biosynthesis-of-secondary-metabolites | 1 | 1 | -1 | 1 | 1 | 1 |
| PROtm-reverse | M-pro–L-m | M-TM-pro–L-c M-pro–L-c | NA | 1 | -1 | 1 | -1 | -1 | 1 |
| PSERS-SC-reverse | M-cmp-c M-h-c M-ps-SC-c | M-TM-cmp-c M-TM-ser–L-c M-cdpdag-SC-c M-ser–L-c | sce00260-Glycine,-serine-and-threonine-metabolism sce00564-Glycerophospholipid-metabolism sce01100-Metabolic-pathways sce01110-Biosynthesis-of-secondary-metabolites | -1 | 1 | 1 | 1 | 1 | 1 |
| PSERSm-SC-reverse | M-cmp-m M-h-m M-ps-SC-m | M-cdpdag-SC-m M-ser–L-m | sce00260-Glycine,-serine-and-threonine-metabolism sce00564-Glycerophospholipid-metabolism sce01100-Metabolic-pathways sce01110-Biosynthesis-of-secondary-metabolites | -1 | 1 | -1 | 1 | 0 | 1 |
| PTD1INOtn-SC-reverse | M-ptd1ino-SC-n | M-ptd1ino-SC-c | NA | -1 | 1 | 1 | 1 | 1 | 1 |

| Reaction data | | | | Wild Type | | | $mip6\Delta$ | | |
|---|---|---|---|---|---|---|---|---|---|
| Rxn | Reactants | Products | Pathways | t0 | t20 | t120 | t0 | t20 | t120 |
| PUNP6-reverse | M-2dr1p-c M-hxan-c | M-din-c M-pi-c | sce00230-Purine-metabolism sce00240-Pyrimidine-metabolism sce00760-Nicotinate-and-nicotinamide-metabolism sce01100-Metabolic-pathways sce01110-Biosynthesis-of-secondary-metabolites | -1 | 1 | 1 | 1 | 1 | 1 |
| SACCD2-reverse | M-akg-c M-h-c M-lys–L-c M-nadh-c | M-TM-lys–L-c M-TM-nad-c M-h2o-c M-nad-c M-saccrp–L-c | sce00300-Lysine-biosynthesis sce00310-Lysine-degradation sce01100-Metabolic-pathways sce01110-Biosynthesis-of-secondary-metabolites sce01230-Biosynthesis-of-amino-acids | -1 | -1 | -1 | 1 | 1 | 1 |
| SERt2r-reverse | M-h-c M-ser–L-c | M-TM-ser–L-c M-h-e M-ser–L-e | NA | 1 | -1 | -1 | 1 | 1 | 1 |
| SUCD2-u6m-reverse | M-fum-m M-q6h2-m | M-q6-m M-succ-m | sce00020-Citrate-cycle-(TCA-cycle) sce00190-Oxidative-phosphorylation sce01100-Metabolic-pathways sce01110-Biosynthesis-of-secondary-metabolites sce01200-Carbon-metabolism | 1 | -1 | 1 | 1 | 1 | 1 |
| THIORDXp-reverse | M-h2o-x M-trdox-x | M-h2o2-x M-trdrd-x | sce04122-Sulfur-relay-system | -1 | -1 | -1 | 1 | 1 | 1 |
| THRt2r-reverse | M-h-c M-thr–L-c | M-h-e M-thr–L-e | NA | -1 | -1 | -1 | 1 | 1 | 1 |
| TRDOXtp-reverse | M-trdox-x | M-trdox-c | NA | 1 | 1 | -1 | 1 | 0 | 1 |
| TRDRDtp-reverse | M-trdrd-x | M-trdrd-c | NA | 1 | 1 | -1 | 1 | 0 | 1 |
| TREt2v-reverse | M-h-v M-tre-v | M-TM-tre-c M-h-c M-tre-c | NA | -1 | 1 | -1 | 1 | 1 | 1 |
| URIDK2r-reverse | M-adp-c M-dudp-c | M-TM-atp-c M-atp-c M-dump-c | sce00240-Pyrimidine-metabolism sce01100-Metabolic-pathways | -1 | -1 | 1 | 1 | 1 | 1 |

##### 6.3 Supplementary Table 2

Genes involved in differential reactions identified by MAMBA and showing a significant increasing mip6 affinity after heat-shock according to Martin-Exposito et al., 2019 [20].

| Gene Name | Description |
|---|---|
| AGP1 | Low-affinity amino acid permease with broad substrate range; involved in uptake of asparagine, glutamine, and other amino acids; expression regulated by SPS plasma membrane amino acid sensor system (Ssy1p-Ptr3p-Ssy5p); AGP1 has a paralog, GNP1, that arose from the whole genome duplication |
| SFA1 | Bifunctional alcohol dehydrogenase and formaldehyde dehydrogenase; formaldehyde dehydrogenase activity is glutathione-dependent; functions in formaldehyde detoxification and formation of long chain and complex alcohols, regulated by Hog1p-Sko1p; protein abundance increases in response to DNA replication stress |
| PDE1 | Low-affinity cyclic AMP phosphodiesterase; controls glucose and intracellular acidification-induced cAMP signaling, target of the cAMP-protein kinase A (PKA) pathway; glucose induces transcription and inhibits translation |
| GRE3 | Aldose reductase; involved in methylglyoxal, d-xylose, arabinose, and galactose metabolism; stress induced (osmotic, ionic, oxidative, heat-shock, starvation and heavy metals); regulated by the HOG pathway; protein abundance increases in response to DNA replication stress |
| TPO1 | Polyamine transporter of the major facilitator superfamily; member of the 12-spanner drug:H(+) antiporter DHA1 family; recognizes spermine, putrescine, and spermidine; catalyzes uptake of polyamines at alkaline pH and excretion at acidic pH; during oxidative stress exports spermine, spermidine from the cell, which controls timing of expression of stress-responsive genes; phosphorylation enhances activity and sorting to the plasma membrane |
| ADH2 | Glucose-repressible alcohol dehydrogenase II; catalyzes the conversion of ethanol to acetaldehyde; involved in the production of certain carboxylate esters; regulated by ADR1 |
| ADH1 | Alcohol dehydrogenase; fermentative isozyme active as homo- or heterotetramers; required for the reduction of acetaldehyde to ethanol, the last step in the glycolytic pathway; ADH1 has a paralog, ADH5, that arose from the whole genome duplication |
| GRE2 | 3-methylbutanal reductase and NADPH-dependent methylglyoxal reductase; stress induced (osmotic, ionic, oxidative, heat-shock and heavy metals); regulated by the HOG pathway; restores resistance to glycolaldehyde by coupling reduction of glycolaldehyde to ethylene glycol and oxidation of NADPH to NADP+; protein abundance increases in response to DNA replication stress; methylglyoxal reductase (NADPH-dependent) is also known as D-lactaldehyde dehydrogenase |
| TPO4 | Polyamine transporter of the major facilitator superfamily; member of the 12-spanner drug:H(+) antiporter DHA1 family; recognizes spermine, putrescine, and spermidine; localizes to the plasma membrane |

##### 6.4 Supplementary Table 3

PEA analysis with all gene expression and metabolomics data.

| Pathway | Combined p-value |
|---|---:|
| Protein processing in endoplasmic reticulum | 0.000 |
| Proteasome | 0.000 |
| Starch and sucrose metabolism | 0.001 |
| Glycolysis / Gluconeogenesis | 0.001 |
| Biosynthesis of secondary metabolites | 0.002 |
| Pentose phosphate pathway | 0.005 |
| Terpenoid backbone biosynthesis | 0.012 |
| Base excision repair | 0.012 |
| Homologous recombination | 0.016 |
| Longevity regulating pathway - multiple species | 0.016 |
| Carbon metabolism | 0.016 |
| Sulfur metabolism | 0.018 |
| Ether lipid metabolism | 0.025 |
| Glycerophospholipid metabolism | 0.028 |
| DNA replication | 0.045 |
